# A comparative epigenome analysis of gammaherpesviruses suggests cis-acting sequence features as critical mediators of rapid polycomb recruitment

**DOI:** 10.1101/639898

**Authors:** Thomas Günther, Jacqueline Fröhlich, Christina Herrde, Shinji Ohno, Lia Burkhardt, Heiko Adler, Adam Grundhoff

## Abstract

Latent Kaposi sarcoma-associated herpesvirus (KSHV) genomes rapidly acquire distinct patterns of the activating histone modification H3K4-me3 as well as repressive H3K27-me3 marks, a modification linked to transcriptional silencing by polycomb repressive complexes (PRC). Interestingly, PRCs have recently been reported to restrict viral gene expression in a number of other viral systems, suggesting they may play a broader role in controlling viral chromatin. If so, it is an intriguing possibility that latency establishment may result from viral subversion of polycomb-mediated host responses to exogenous DNA.

To investigate such scenarios we sought to establish whether rapid repression by PRC constitutes a general hallmark of herpesvirus latency. For this purpose, we performed a comparative epigenome analysis of KSHV and the related murine gammaherpesvirus 68 (MHV-68). We demonstrate that, while latently replicating MHV-68 genomes readily acquire distinct patterns of activation-associated histone modifications upon *de novo* infection, they fundamentally differ in their ability to efficiently attract H3K27-me3 marks. Statistical analyses of ChIP-seq data from *in vitro* infected cells as well as *in vivo* latency reservoirs furthermore suggest that, whereas KSHV rapidly attracts PRCs in a genome-wide manner, H3K27-me3 acquisition by MHV-68 genomes may require spreading from initial seed sites to which PRC are recruited as the result of an inefficient or stochastic recruitment, and that immune pressure may be needed to select for latency pools harboring PRC-silenced episomes *in vivo*.

Using co-infection experiments and recombinant viruses, we also show that KSHV’S ability to rapidly and efficiently acquire H3K27-me3 marks does not depend on the host cell environment or unique properties of the KSHV-encoded LANA protein, but rather requires specific cis-acting sequence features. We show that the non-canonical PRC1.1 component KDM2B, a factor which binds to unmethylated CpG motifs, is efficiently recruited to KSHV genomes, indicating that CpG island characteristics may constitute these features. In accord with the fact that, compared to MHV-68, KSHV genomes exhibit a fundamentally higher density of CpG motifs, we furthermore demonstrate efficient acquisition of H2AK119-ub by KSHV and H3K36-me2 by MHV-68 (but not vice versa), furthermore supporting the notion that KSHV genomes rapidly attract PRC1.1 complexes in a genome-wide fashion. Collectively, our results suggest that rapid PRC silencing is not a universal feature of viral latency, but that some viruses may rather have adopted distinct genomic features to specifically exploit default host pathways that repress epigenetically naive, CpG-rich DNA.

**Author Summary:** During herpesvirus latency, viral genomes persists as partially repressed nuclear episomes which do not express genes required for progeny production. Latently infected cells not only form a reservoir of lifelong persistence but also represent the driving force in cancers associated with tumorigenic herpesviruses such as KSHV. Hence, it is fundamentally important to understand the mechanisms controlling latency. We have shown previously that latent KSHV episomes rapidly acquire H3K27-me3, a histone mark associated with polycomb repressive complexes (PRC). PRCs play a pivotal role in the control of developmental genes but are also involved in the pathogenesis of several tumors. We here investigated whether PRC-repression represents a general feature of herpesvirus latency. By performing side-by-side analyses of KSHV and the related MHV-68 we show that the latter indeed has a fundamentally lower propensity to acquire H3K27-me3, and that KSHV’S ability to rapidly attract this mark is most likely the result of a specific sequence composition that promotes recruitment of non-canonical PRC1 (a complex which is important for the regulation of cellular CpG islands). Our results have widespread implications for nuclear DNA viruses and suggest that some viruses have specifically evolved to exploit common host responses to epigenetically naive DNA.

## Introduction

Herpesvirus latency is characterized by nuclear persistence of viral episomes in the absence of viral progeny production. To establish latency, herpesviruses must ensure that genes required for productive/lytic infection are efficiently silenced, while expression of those required for episomal persistence must be preserved. Given these requirements, epigenetic modifications of viral DNA or chromatin have long been suspected to play an important role during establishment and maintenance of herpesvirus latency.

Kaposi Sarcoma-associated herpesvirus (KSHV) is the etiologic agent of several tumors, including Kaposi sarcoma (KS), primary effusion lymphoma (PEL) and the plasmablastic variant of multicentric Castleman disease (MCD) [reviewed in 1, 2]. The tumor cells in these malignancies unequivocally express the major latency cassette of KSHV to produce several viral microRNAs (miRNAs) and at least four proteins. Among the latter, the multifunctional latency-associated nuclear antigen (LANA, encoded by ORF73) is especially important for viral persistence: LANA recruits the host cell replication machinery to viral origins of replication and tethers KSHV genomes to host chromosomes to ensure proper episome segregation [reviewed in 3]. While latent KSHV transcription and replication have first been studied in B cell lines derived from PEL tumors, later studies have shown that the virus readily establishes latency in a wide variety of adherent cells [4, 5]. Using such models, we have previously shown that methylation of viral DNA is a late and secondary event that is preceded by the establishment of distinct histone modification patterns [6]. Most notable among these are the global acquisition of the repressive histone mark H3K27-me3 (a modification deposited by the EZH2 component of polycomb repressive complex 2, PRC2) and formation of distinct peaks of activation-associated histone marks (H3K4-me3 and H3K9/K15-ac) not only at the major latency promoter, but also various regions encoding transcriptionally silent lytic genes. While the latter are generally anti-correlated with H3K27-me3, they co-exist with the repressive mark on a few loci (including the promoter of the major lytic transactivator Rta, encoded by ORF50) to form bivalent chromatin, a type of facultative heterochromatin poised for rapid re-expression. The putative function of the remaining activation-associated histone marks in transcriptionally silent latent chromatin, however, remains unknown. Later studies have confirmed and extended above findings to demonstrate, for example, that acquisition of activating and repressive histone marks represent temporally separate processes, with the former already being fully present at 24h post infection, whereas H3K27-me3 marks require approximately 3 days to reach peak levels [7-9]. H3K27-me3 patterns furthermore evolve uniformly over the entire genome, sparing only some of the regions which already carry activation-associated modifications [8]. This observation strongly suggests that KSHV episomes attract PRCs via a global mechanism, rather than spreading from initial sites to which they are recruited by sequence-specific transcription factors (as is likely the case for activation-associated marks). In addition to de novo infected cells and tumor-derived cell lines, widespread H3K27-me3 marks have also been found to decorate KSHV genomes in primary KS tumors [10]. The fact that knock-down or pharmacological inhibition of PRC2 or PRC1 results in an increase of lytically reactivating cells furthermore supports the hypothesis that polycomb repression plays an important role in the maintenance of KSHV latency [6, 7, 11, 12].

An important question that remains, however, is what cellular mechanisms shape the viral epigenome and to what extent such mechanisms, in particular PRC recruitment, are influenced by viral factors. Indeed, although a recent study by Toth and colleagues elegantly demonstrated a principal requirement for LANA during H3K27-me3 acquisition, the kinetics of H3K27-me3 accumulation and LANA expression suggest that other, as of yet unknown factors are critically involved in the initial global recruitment process [13]. Since polycomb group proteins (PcG) have recently emerged as important factors that restrict viral gene expression of diverse viral species including CMV, HSV-1 and HIV, it is an intriguing possibility that polycomb repression and H3K27-me3 deposition may represent a default host response to the nuclear presence of non-chromatinized and/or unmethylated DNA. In such scenarios, latency establishment may result from viral subversion of host cell mechanisms which serve to silence epigenetically naive DNA. Apart from KSHV, however, the chromatin landscape associated with the early phase of herpesvirus latency remains unknown.

This is also true for murine gammaherpesvirus-68 (MHV-68), a virus which shares a co-linear genome organization as well as significant functional and sequence conservation across a large fraction of its protein products with KSHV. Since MHV-68 is furthermore able to establish persistent infections in laboratory mice it represents a valuable model system to study infection by KSHV and related γ-herpesviruses. Similar to KSHV, MHV-68 persists in CD19+ B-cells, with germinal center and marginal zone B-cells exhibiting the highest frequency of latent infection [14]. While the virus can induce lymphoproliferative disease under certain experimental conditions, the clonal lymphoma cells are typically MHV-68-negative, suggesting tumor induction via hit-and-run mechanisms, paracrine effects or chronic inflammation [15]. Indeed, to date only a single persistently infected B cell line (S11) has been established from infected mice [16]. Since (unlike KSHV) MHV-68 by default enters the lytic cycle upon *in vitro* infection of permissive cells, S11 and its derivative sub-clones continue to serve as the predominant tissue culture model to study MHV-68 latency.

Epigenetic analyses of latent MHV-68 episomes thus far has been limited to the ORF50 promoter region. Although this region is generally poor in CpG dinucleotides, Gray and colleagues found that the distal ORF50 promoter undergoes CpG-methylation at later stages of latency in mice [17]. In accord with this, Yang et al. found the ORF50 promoter to be methylated in mouse derived splenocytes as well as in S11E (a sub-clone of S11 which shows a lower rate of spontaneous reactivation) cells [18]. However, treatment with the DNA-methyltransferase inhibitor 5’-Azacytidine failed to induce lytic replication, suggesting that DNA methylation is not the primary or only block of ORF50 expression during latent MHV-68 infection.

Given the above, we set out to perform a comparative and comprehensive analysis of epigenetic modifications in latent KSHV and MHV-68 genomes. In particular, we aimed at elucidating whether latently replicating MHV-68 episomes would attract polycomb repressive complexes (PRCs) with similar efficiency as KSHV, thus arguing for PRC recruitment representing a general characteristic of gammaherpesvirus latency. Indeed, our results show that, even in the same host background, MHV-68 genomes acquire H3K27-me3 marks much slower and less efficiently than KSHV, suggesting that virus specific cis-acting features, most likely the presence of a high frequency of unmethylated CpG motifs, regulate the recruitment of PRCs to epigenetically naïve viral DNA.

## Results

### Comparative analysis of gammaherpesvirus epigenomes in tumor-derived B-cell lines

Although MHV-68 establishes latency in the spleen of infected mice, the low frequency of such cells against a high background of uninfected cells greatly complicates direct ChIP-seq analysis of primary splenic lymphocytes. We therefore sought to first investigate the epigenetic landscape of MHV-68 genomes in the established B-cell line S11E and compare it to that of KSHV in BCBL-1 cells, a cell line derived from primary effusion lymphoma. As shown in Fig 1A, our ChIP-Seq analysis of BCBL1 cells produced profiles of activating H3K4-me3 and repressive H3K27-me3 marks which were highly similar to those of previous ChIP on microarray analyses [6, 11]. In contrast, ChIP-seq profiles of MHV-68 genomes in S11E cells were markedly different (see Fig 1B). Although, like KSHV, MHV-68 epigenomes exhibited peaks of activation-associated histone marks outside of the latency region, the total number as well as relative enrichment of such peaks was substantially lower. Overall, MACS14 peak detection found only three regions to be significantly enriched for H3K4-me3: two peaks upstream of M2 and M7 and another peak in the 3’ region of ORF75A (Fig 1B, black bars). Two additional sites upstream of ORF18b and M11 (marked with an asterisk in Fig 1B) were also enriched in the IgG control (Fig S1A) and thus considered false positives.

**Fig. 1:**
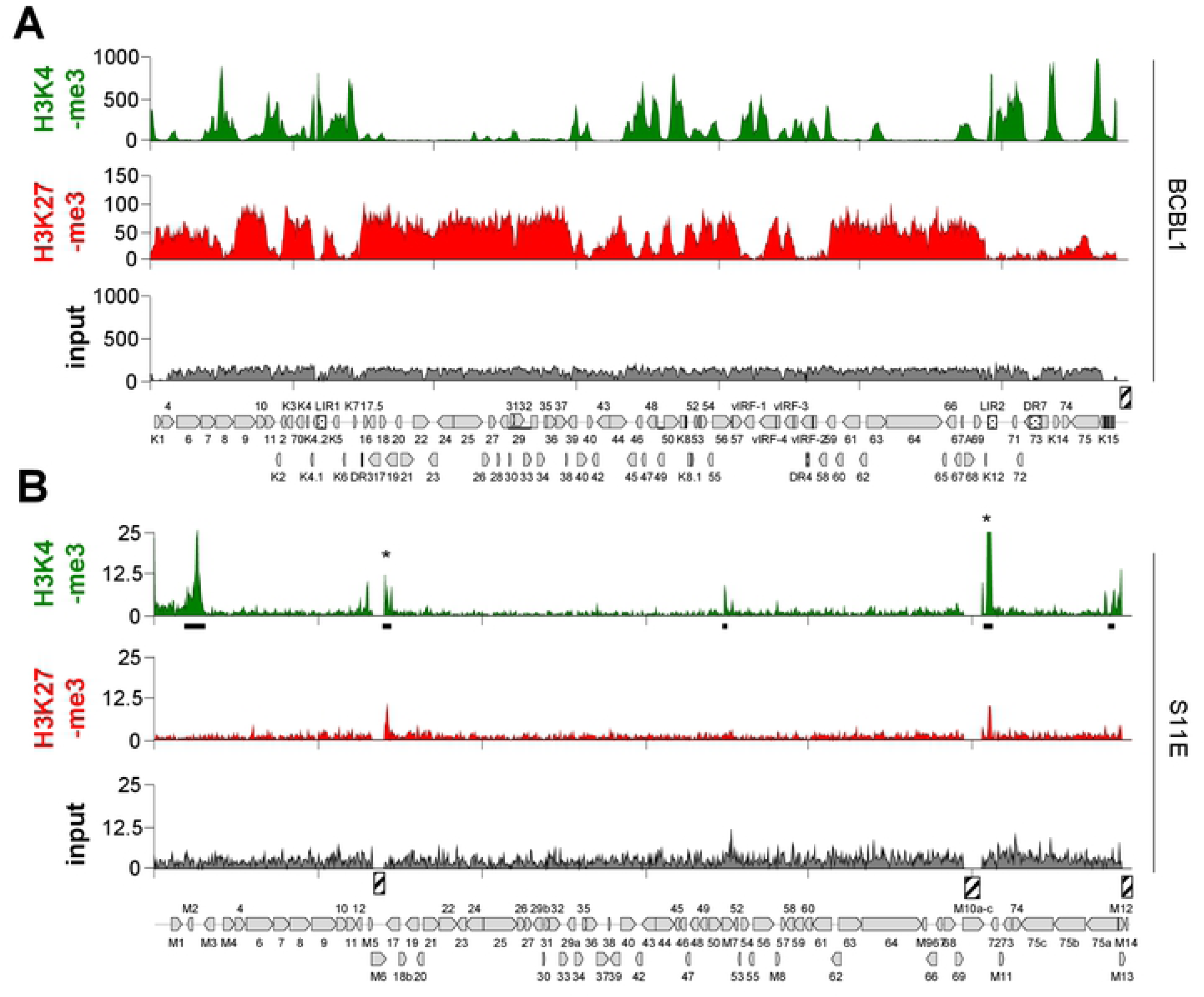
ChIP-seq analysis of MHV-68 and KSHV epigenomes in tumor-derived B-cell lines. ChiP-seq coverage for H3K4-me3 and H3K27-me3, or corresponding input coverage across the KSHV genome in BCBL1 (A) or the MHV-68 genome in S11E cells (B) for the indicated antibodies. Regions on the MHV-68 genome enriched for H3K4-me3 as detected using MACS14 are indicated by black bars. Asterisks indicate likely false positives also present in the IgG control (S1A Fig). Hashed boxes indicate repetitive regions (including the terminal repeats) that were excluded from the analysis as they do not allow unique read mapping.

In stark contrast to KSHV, we did not detect a distinct profile or abundant levels of the facultative heterochromatin mark H3K27-me3 on MHV-68 episomes in S11E cells. The constitutive heterochromatin mark H3K9-me3 was also absent (see S1 Fig A for H3K9-me3 and H3K9/K14-ac profiles). To further investigate whether the weak signals of repressive histone marks may nevertheless reflect a biologically significant modification profile we calculated correlation coefficients (cc) for the different data sets (see S1 and S2 Dataset for coverage data and pairwise correlation coefficients, respectively, across all ChIP-seq experiments in this study). Since H3K4-me3 and H3K27-me3 are (except at bivalent loci) generally anti-correlated, the absence of such a correlation would indicate that the H3K27-me3 profiles represent pure background. Indeed, consistent with our previous findings [6], KSHV showed profound anti-correlation between H3K4-me3 and H3K27-me3 (cc: −0.55). In contrast, our MHV-68 data show no anti-correlation of both histone marks (cc: 0.08) indicating that the signal of H3K27-me3 is not distinguishable from the background ChIP signal. Hence, our analysis suggests that the majority of MHV-68 episomes in S11E cells do not carry significant levels of repressive histone marks and appear to be decorated by only a small number of activation associated peaks.

### RNA-seq analysis suggests that lytic gene products dominate the viral transcriptome in S11E cells

In KSHV-positive cells, reactivation of productive replication does result in only minor alterations of the bulk epigenetic profile. During late stages of the lytic cycle, accumulation of large numbers of *de novo* replicated genomes (which do not carry any histones and thus only contribute to the input) furthermore decreases relative ChIP enrichment values. We therefore wondered whether the sparsity of detectable ChIP signals as shown in Fig 1B may, at least in part, result from spontaneously reactivating cells carrying a disproportionally large fraction of epigenetically naive episomes which mask the authentic epigenetic profile of latent episomes. Although a predominantly latent expression profile of the original S11E had been shown by array hybridization [19], we wished to compare the transcription profile of our S11E subculture to lytic MHV-68 transcription during the course of a *de novo* infection. For this purpose, we performed RNA-seq of our S11E cells and murine cultures of the epithelial cell line MLE12 undergoing lytic replication after *de novo* infection with MHV-68 (Fig 2, see S2 Fig for additional time points post infection).

**Fig. 2:**
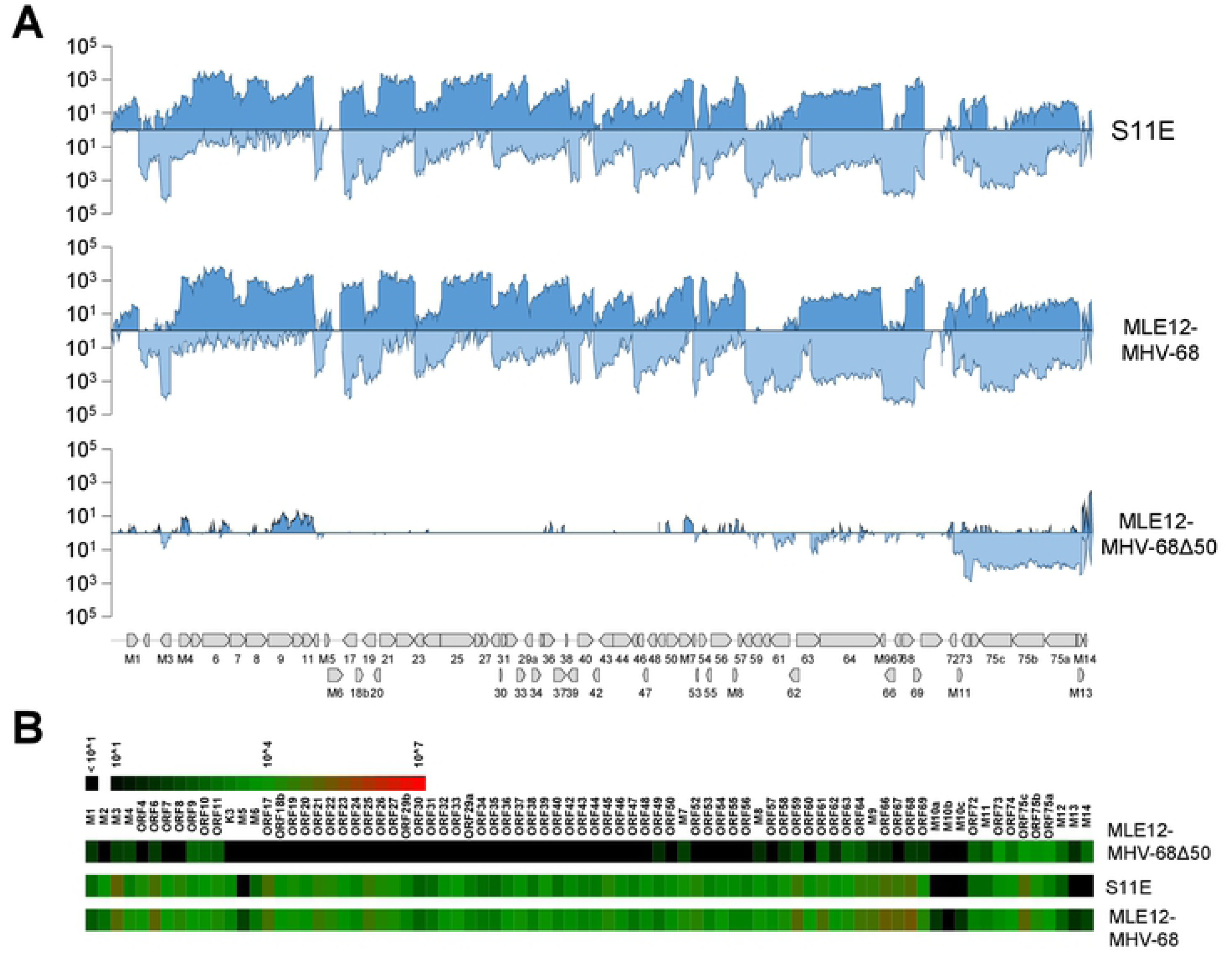
RNA-seq analysis of S11E and de novo infected MLE12 cells. (A) RNA-seq analysis of persistently infected S11E cells (upper panel), *de novo* MHV-68 infected MLE12 cells at 12 hours post infection (center panel) or GFP-sorted MLE12 cells which had been infected with MHV-68Δ50 for more than 3 weeks (lower panel). RNA sequencing was performed using a strand-specific sequencing protocol and resulting paired-end RNA-seq reads were mapped to the MHV-68 reference sequence (NC_001826) using the splice-sensitive STAR pipeline (see Material and Methods for details). Coverage tracks depict mean coverage across 100 bp binning windows. Forward and reverse strand coverage is shown in the upper and lower plots of each panel. (B) Heatmaps depicting normalized read coverage across individual MHV-68 ORFs annotated in the NC_001826 GenBank entry for the experiments shown in A.

Surprisingly, the bulk transcription profile of our S11E cultures was highly similar to that of the lytically infected MLE12 cells, indicating that the transcriptional activity of viral episomes in spontaneously reactivating cells greatly outweighs that of the numerically more abundant latently infected cells. Although we cannot directly determine the ratio of productively and latently replicating episomes in this population, these data indicate that the epigenetic profiles observed in the S11E cells may only partially reflect the authentic chromatin state of latent MHV-68 genomes. The fact that H3K27me3 profiles of latent KSHV genomes remain clearly detectable even in BCBL1 cultures that are induced by TPA [11], however, argues for the assumption that the latent fraction of MHV-68 genomes in S11E cells does indeed not carry abundant levels of this mark.

### An ORF50 deletion mutant (MHV-68Δ50) establishes a strictly latent gene expression profile

To circumvent the experimental and interpretational limitations imposed by S11-derived cultures, we sought to establish a strictly latent *de novo* infection system. For this purpose, we generated a mutant MHV-68 clone (MHV-68Δ50) with a deletion encompassing the region encoding Rta, the master lytic transactivator (see methods section for information on generation of the mutant and production of infectious viral particles). We reasoned that, due to its inability to enter the lytic cycle, the mutant should allow us to observe the epigenetic profile acquired by latently replicating episomes, in particular with regard to the hallmark traits of latent KSHV genomes, namely activation-associated marks outside of latent transcription units and global acquisition of repressive H3K27-me3 marks.

Infection of murine lung epithelial MLE12 cells with MHV-68Δ50 resulted in fully viable cultures which, similar to *de novo* latently infected KSHV cell lines [4], spontaneously lost viral episomes at an estimated rate of approx. 5-10% per cell generation. We maintained a highly positive culture by repeated sorting for GFP-positive cells over a period of more than three weeks and subsequently analyzed the transcriptome in these long-term infected cells by strand-specific RNA-seq. As shown in Fig 2B and -C (see also S3 Dataset), in contrast to our S11E cells MHV-68Δ50 infected cells indeed exhibited a viral gene expression profile that was highly restricted. The highest level of viral gene expression could be assigned to ORF73, an ORF which encodes the MHV-68 homologue of LANA (mLANA). Additionally, abundant coverage was detected in the region encompassing ORFs 75a to 75c, likely due to the presence of unspliced variants of ORF73-coding transcripts that originate at the terminal repeats. Notably, a weak (note logarithmic scale in Fig 2B) but distinct transcription signature was detectable at two broader sites of the genome between ORFs M1 and 11 as well as ORFs 49 and 69.

### Epigenetic analysis of MHV-68Δ50 reveals distinct profiles of activation-associated histone marks and suggests that viral genomes do not efficiently recruit PRCs

Given the highly restricted transcription patterns described above, we concluded that MHV-68Δ50 infected cells provide a uniform population of latently replicating episomes and proceeded to analyze the distribution of activating and repressive histone modifications by ChIP-seq. In parallel, we performed ChIP-seq from long-term *in vitro* infected SLK cells (SLK_P_) [4] to allow direct comparison with a strictly latent KSHV infection. Similar to BCBL1, the ChIP-seq results for SLK_P_ cells were highly congruent with our previous ChIP on microarray data [6], exhibiting profound anti-correlation (cc: - 0.58) of strongly enriched H3K4-me3 and H3K27-me3 profiles (Fig 3A). Hence, as noted before [6, 8], while a few H3K4-m3 peaks differ between BCBL1 and SLK_P_ cells (likely due to differences in the cell type-specific transcription factor repertoire), all regions not already decorated by H3K4-me3 acquire abundant H3K27-me3 marks, indicating that global affinity for PRCs is a fundamental property of latent KSHV episomes.

**Fig. 3:**
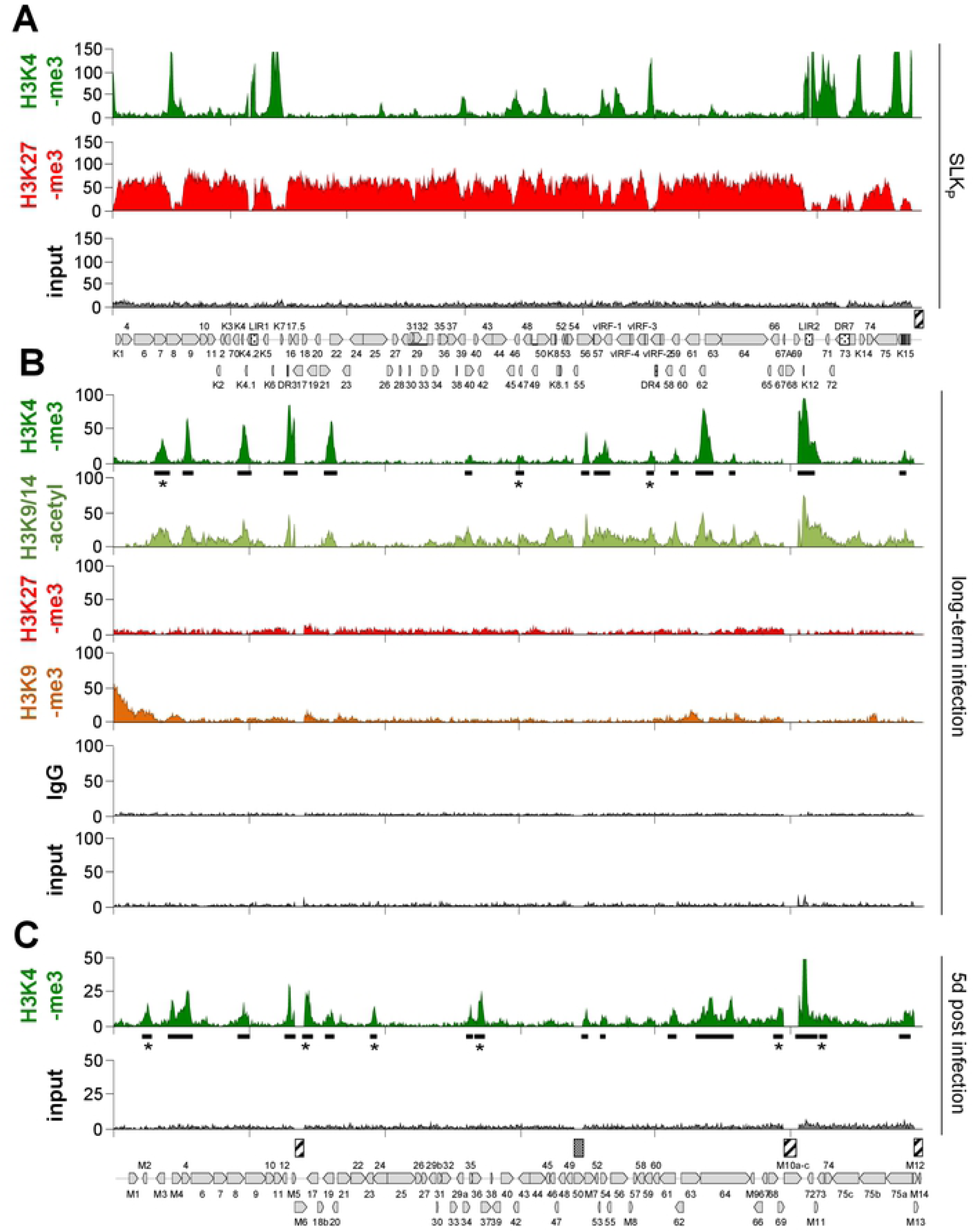
ChIP-seq analysis of KSHV and MHV-68Δ50 epigenomes in SLKp or MLE12 cells. ChIP-seq results from (A) KSHV infected SLK_P_ cells or (B) long-term MHV-68Δ50 infected MLE12 cells and (C) MLE12 cells at day 5 post infection with MHV-68Δ50. Solid bars underneath H3K4-me3 plots indicate peak regions as detected by MACS14. Regions that were uniquely enriched at either day five or in long-term latency are marked with an asterisk (*) in each panel. Hashed boxes above the MHV-68 and KSHV genomes map indicate repetitive regions excluded from the analysis, and the dotted box above MHV-68 indicates the deletion of ORF50 coding sequences in MHV-68Δ50.

As shown in Fig 3B, MHV-68Δ50 indeed also acquired a highly distinct profile of activating histone marks (H3K4-me3 and H3K9/K14-ac), with MACS14 peak detection revealing 15 significantly enriched H3K4-me3 positive sites (black bars in Fig 3B). Interestingly, while latently replicating MHV-68Δ50 and KSHV episomes were similar in their recruitment of activation-associated histone marks, MHV-68Δ50 did not attract abundant levels of H3K27-me3. Nevertheless, despite overall low enrichment the resulting profiles showed anti-correlation with H3K4-me3 and H3K9/K14-ac (cc: −0.25 and −0.34, respectively), indicating a baseline signature of H3K27-me3 that may result from slow recruitment of PRCs by a subset of viral episomes. The constitutive heterochromatin mark H3K9-me3 was likewise not widely distributed throughout the MHV-68Δ50 genome, but also showed slight anti-correlation with H3K9/K14-ac (cc: −0.25). While a region encompassing ORFs M1 to M4 at the left end of the genome was noticeably enriched for H3K9-me3 marks, mapping of the ChIP-seq reads to the BAC (which is inserted upstream of M1) also revealed profound H3K9-me3 enrichment along the integrated cassette (S1 Fig B). Hence, at least a subset of the BAC sequences undergo heterochromatinization, and it is thus possible that the observed signals in the M1-M4 region result from spreading of such heterochromatin into the neighboring viral regions.

To further elucidate the timeframe during which activation-associated histone mark patterns are established we infected MLE12 cells with MHV-68Δ50 and analyzed episomes at day five post infection by ChIP-Seq (Fig 3C). Indeed, the H3K4-me3 patterns observed at this early time point were highly similar (average cc: 0.55), but not completely identical to those detected after long-term infection. ChIP-peak calling using MACS14 on the H3K4-me3 samples detected a total of 9 uniquely enriched sites (marked by asterisks in Fig 3B and C) located upstream of M2, ORF18b, ORF23, ORF37, ORF69/M10 ORF73/74 in long-term infected cells, or upstream of M3/M4, ORF42 and ORF59 at day five post infection (see S4 Dataset for a list of all H3K4-me3 peaks identified during our experiments). Interestingly, although we did not experimentally interrogate H3K4me3-marked regions for promoter activity, cross-comparison of peak location with transcription cassettes predicted from our RNA-seq data (see S1 Protocol, S5 Dataset) suggests that these peaks are preferentially found upstream of genes expressed immediately after *de novo* infection or reactivation (S3 Fig).

### Statistical analysis of ChIP-seq data results confirms overall absence of H3K27-me3 from MHV-68Δ50 genomes

Although we did not observe that MHV-68Δ50 genomes were strongly enriched for H3K27-me3 (Fig 3), the anti-correlation with H3K4-me3 suggests that the weak H3K27-me3 signals did not purely represent background. We therefore set out to perform a statistical analysis designed to provide a better estimate of the actual H3K27-me3 levels acquired by MHV-68 and KSHV episomes via direct comparison with strongly enriched host regions. For this purpose, we mapped the ChIP-seq and input reads from two independent experiments of long-term MHV-68Δ50-infected MLE12 or KSHV-infected SLK_P_ cells to the appropriate host genome assemblies and, using enrichment detection with SICER/EPIC [20], identified the 200 host regions most significantly enriched for H3K27-me3. Conversely, as a negative control we randomly selected 200 regions with a similar length distribution from the host genome, excluding all regions previously identified as H3K27-me3 enriched by SICER/EPIC2 as well as excluding regions blacklisted according to ENCODE. Viral H3K27-me3 enrichment values were subsequently calculated by sliding a window of 10 kb (to reflect broad regions comparable to detected host regions) across the viral genome. To allow cross-comparison between individual samples, all data were then normalized to the median value of the negative control regions. The data shown in Fig 4 thus represent fold enrichment over the negative control loci (cellular ChIP background) and allow a direct assessment of H3K27-me3 occupancy on viral genomes in the overall context of the host epigenome (note that this is an assessment of global H3K27-me3 acquisition across the viral genome, rather than recruitment to individual and discrete sites). For more details, see the materials and methods section. As shown in the right panel of Fig 4, enrichment of H3K27-me3 on KSHV genomes in SLK_P_ cells was significantly higher compared to not only the negative, but even the positive host control loci, underlining our previous notion that efficient and global recruitment of PRCs is a hallmark of KSHV latency. In stark contrast, average H3K27-me3 enrichment levels on MHV-68Δ50 genomes (left panel) were comparable to those of the negative control set. Thus, our statistical analysis strongly supports the conclusion that, even after prolonged *in vitro* infection, the majority of latently replicating MHV-68Δ50 genomes does not efficiently attract H3K27-me3 marks.

**Fig. 4:**
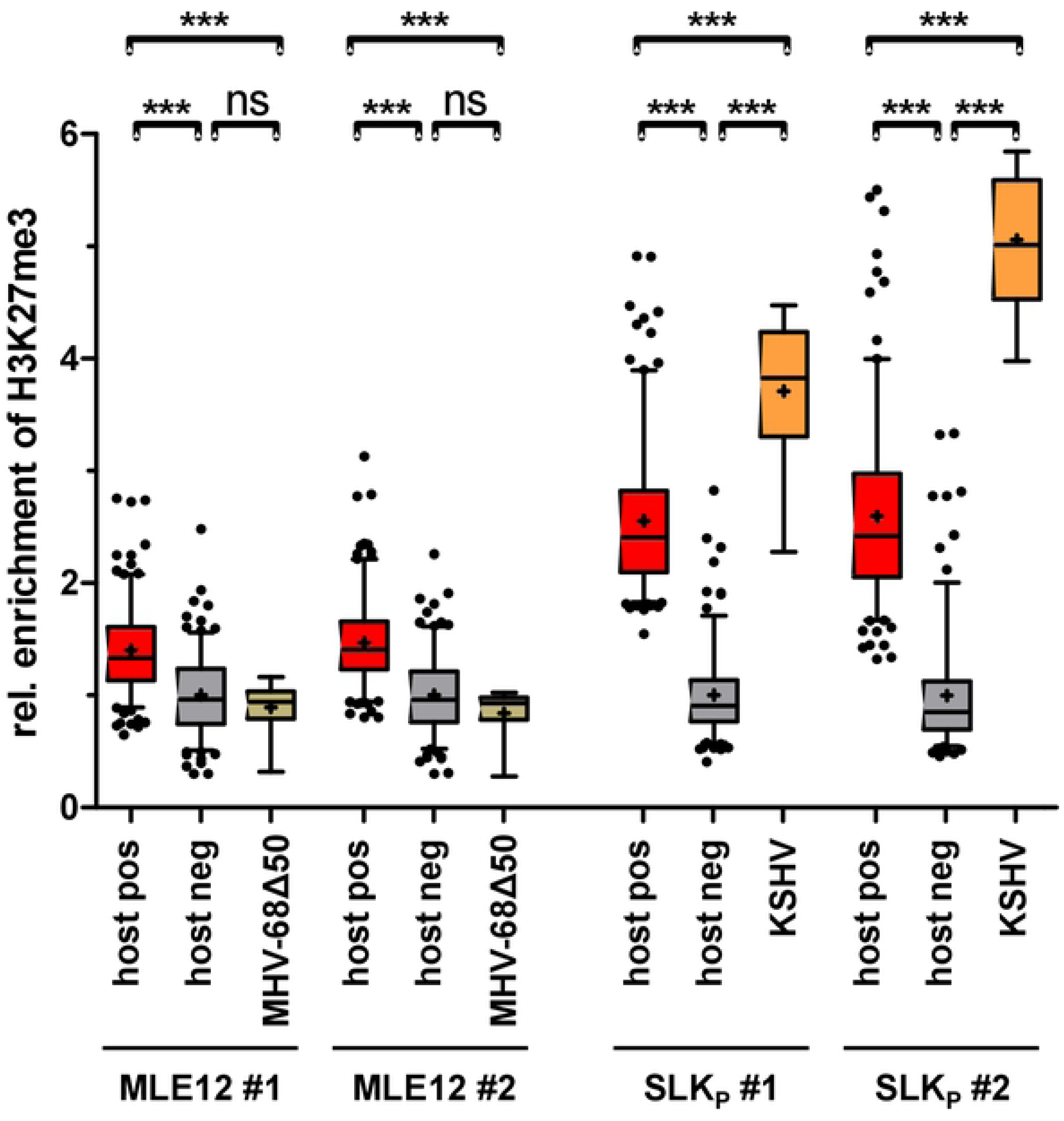
Statistical analysis of H3K27-me3 levels acquired by viral episomes. Shown is an analysis of two biological replicates collected from MHV-68Δ50-infected MLE12 cells (left panels) or KSHV-infected SLKp cells (right panels) each. The graphs depict average enrichment of H3K27-me3 levels in positive and negative control regions of the host genomes, relative to average enrichment across the viral genome. H3K27-me3 positive (host pos) and negative (host neg) host regions (200 each) were detected by SICER/EPIC2, and enrichment across viral sequences was calculated using a 10kb sliding window (see methods section for details). For each region we individually calculated the H3K27-me3 to input read count ratio and normalized all groups to the average of the respective negative control. Data are shown as box-whisker-plots with 5th-95th percentile and median (+). Significance was calculated by 1way ANOVA testing (MLE-12: F = 64.36, df = 817; SLK_P_: F = 341, df = 770). Significance is indicated by asterisks or ns (not significant).

### The ability to rapidly recruit PRC2 is a virus-but not a host cell-specific feature

Since the lack of H3K27-me3 (and thus, most likely, PRC2-) recruitment to MHV-68Δ50 episomes was in striking contrast to the mechanisms observed in latent KSHV infection, we sought to investigate whether this disparate behavior indeed reflected intrinsic properties of the two viruses or may rather be caused by the host cell background (i.e., a potentially general impairment of polycomb recruitment pathways in the murine system). We therefore performed a superinfection of long-term MHV-68Δ50-infected MLE12 cells with KSHV for five days, a timeframe that allows full establishment of latency associated chromatin structures on KSHV episomes after *de novo* infection of human cells [6, 8]. We subsequently analyzed the ability of KSHV to recruit H3K27-me3 marks in the MHV-68Δ50 positive murine cell system by ChIP-seq (Fig 5), normalizing read counts by input for each virus to correct for differences in copy numbers per cell. Although slightly noisier compared to the previous analysis, H3K4-me3 patterns on MHV-68Δ50 genomes were highly similar to the data shown in Fig 3B (cc: 0.85). Likewise, despite the heterologous host species background, KSHV acquired activation mark patterns that overall correlated well with the profiles observed in SLK_P_ cells (cc: 0.76), with the highest enrichment detectable in the latency associated region. A number of prominent peaks observed in SLKp cells (e.g., the peak upstream of vIRF3), however, were less pronounced in MLE12 cells, likely due to differences with which murine transcription factors bind to specific target sites in the KSHV genome. To determine the extent of H3K27-me3 recruitment by KSHV and MHV-68Δ50 genomes, we performed the same statistical analysis as for the previous dataset (Fig 5B). Indeed, while MHV-68Δ50 genomes were again not significantly different from the background, the enrichment of H3K27-me3 on KSHV genomes was highly significant and (similar to the observations made in SLK_P_ cells) exceeded that of the endogenous murine positive controls. As before, global H3K27-me3 patterns were anti-correlated with H3K4-me3 marks (cc: −0.32).

**Fig. 5:**
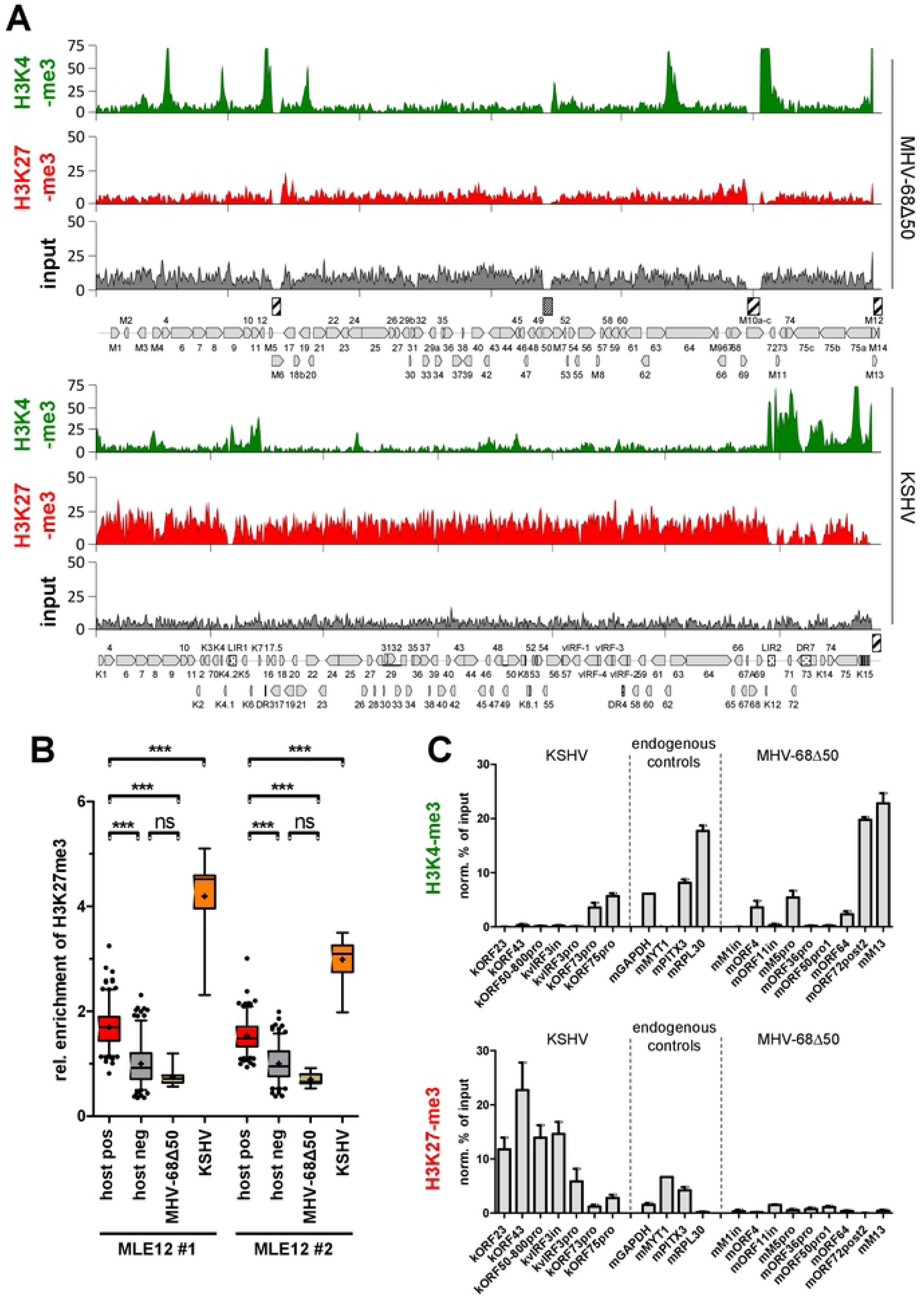
ChIP-seq analysis of KSHV and MHV-68Δ50 epigenomes in superinfected MLE12 cells. (A) H3K4-me3 and H3K27 ChIP-seq coverage across MHV-68Δ50 (top) and KSHV (bottom) genomes in long-term MHV-68Δ50-infected MLE12 cells superinfected with KSHV for five days. Read counts in all samples were normalized to input DNA to correct for differences among KSHV and MHV-68Δ50 episome copy numbers per cell. (B) Statistical analysis of H3K27-me3 enrichment in two replicates of MHV-68Δ50-positive MLE12 cells superinfected with KSHV. Analysis was performed analogous to that shown in Fig 4. Data are shown as box-whisker-plots with 5th-95th percentile and median (+). Significance was calculated by 1way ANOVA testing (F = 270.8, df = 889). Significance is indicated by asterisks or ns (not significant). (C) Confirmatory ChIP-qPCR using KSHV-, MHV-68- and mouse-specific primers as indicated (n=3). Positive controls were as follows: H3K4-me3: GAPDH, RPL30; H3K27-me3:MYT1; H3K4-me3/H3K27-me3 (bivalent chromatin): PITX1. Data are represented as mean ± SEM.

In complete agreement with our ChIP-seq analysis, ChIP-qPCR for a number of viral and host loci confirmed that enrichment of H3K27-me3 on KSHV was about three fold higher when compared to positive control regions, whereas the selected MHV-68Δ50 loci were negative (Fig 5C). Additionally, the ChIP-qPCR confirmed the patterns of activation-associated histone marks in both viruses.

### Unique properties of KSHV LANA are not responsible for rapid H3K27-me3 acquisition

A recent report by Toth et al. [13] demonstrated that acquisition of H3K27-me3 marks by KSHV is critically dependent on the presence of an intact LANA molecule, suggesting that KSHV LANA (kLANA) itself may mediate PRC recruitment. Given this, we wondered whether unique properties of kLANA that are not conserved in murine homolog (mLANA) may be required for rapid acquisition of H3K27-me3, potentially explaining the absence of this mark from MHV-68 episomes in singly or co-infected cells. Although KSHV and MHV-68 LANA preferentially bind to their cognate sites within the terminal repeats, a number of recent studies have suggested that they can also reciprocally support genome replication and maintenance [21-24]. We therefore sought to investigate the ability of a recombinant MHV-68 genome harboring a kLANA gene to attract H3K27-me3 marks. For this purpose, we introduced the ORF50 deletion in the background of a chimeric MHV-68 mutant (recently generated by Habison and colleagues [21]) in which mLANA was replaced by kLANA, permitting us to investigate the chromatin landscape of latently replicating MHV-68 genomes that are maintained by kLANA instead of mLANA. As expected, MLE12 cells infected with either the mLANA-expressing MHV-68Δ50 virus or the MHV-68Δ50-kLANA mutant showed readily detectable expression of the respective LANA protein (Fig 6D). As shown in Figs 6A and B, however, ChIP-seq analysis did not find any significant differences between H3K27-me3 enrichment levels at 5 days post-infection, yielding profiles that were very close to background regardless of whether MHV-68 genomes expressed m- or kLANA proteins. Quantitative ChIP-PCR from chromatin harvested after 5 and 35 days post-infection confirmed these results and demonstrated that, even after prolonged infection, kLANA-mediated episome maintenance did not confer the ability to rapidly attract polycomb repressive complexes to MHV-68 genomes (Fig 6C).

**Fig. 6:**
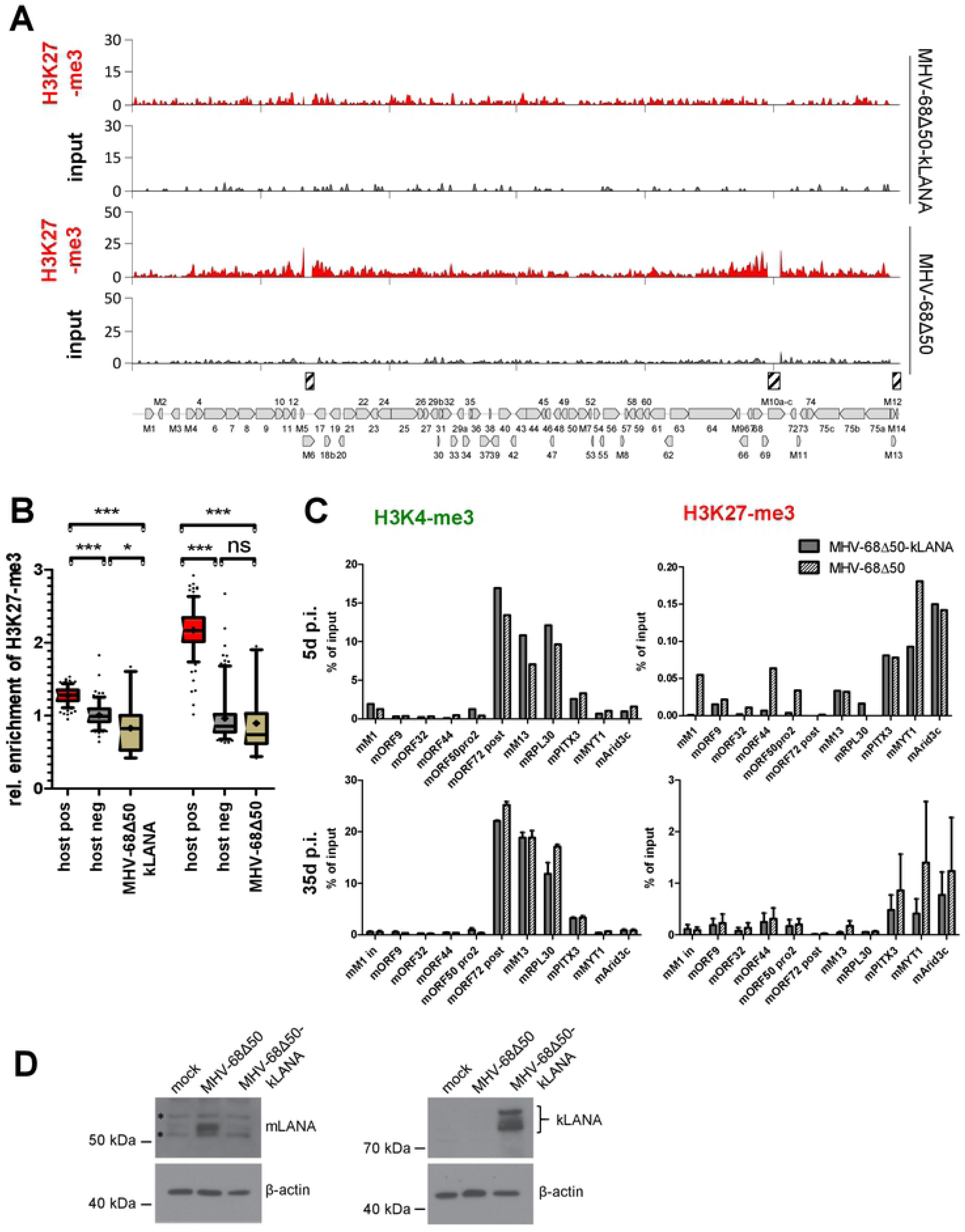
kLANA-expressing MHV-68Δ50 genomes do not gain the ability to rapidly recruit H3K27-me3 marks. (A) H3K27-me3 ChIP-seq coverage across the MHV-68Δ50 genome in MLE12 cells sorted for GFP expression at day 5 post infection with MHV-68Δ50-kLANA (top) or MHV-68Δ50 (bottom). Due to the low titers of the MHV-68Δ50-kLANA virus, ChIP-seq was performed for both cultures using a low-cell ChIP protocol as described in the material and methods section. (B) Relative H3K27-me3 enrichment analysis for the data shown in panel A, performed as described in the legend to Fig 4. Significance was calculated by 1way ANOVA testing (F = 640.7, df = 781) and is indicated by asterisks or ns (not significant). (C) Confirmatory ChIP-qPCR analysis of MHV-68Δ50-kLANA or MHV-68Δ50-infected MLE12 cells at 5 days post infection (n=1, top panel), or after a 35 day period during which cells were repeatedly sorted to achieve a population of 100% GPF positive cells (n=2, bottom panel). Data are represented as mean ± SEM.(D) Western blot analysis of mLANA and kLANA expression in cells infected with MHV-68Δ50-kLANA or MHV-68Δ50, respectively. Unspecific bands are indicated by asterisks.

While kLANA has been shown to support episome maintenance of MHV-68 genomes [21-23], it seemed formally possible that it may be selectively defective for PRC recruitment in a non-syngeneic context, for example due to differential binding stoichiometry at the terminal repeats or the lack of low affinity binding sites in the coding region. We therefore also tested whether trans-complementation of a LANA-deleted KSHV genome with either kLANA or mLANA would result in differential behavior towards PRC repression, expecting that mLANA-maintained KSHV genomes may be unable to acquire H3K27-me3 marks. Accordingly, we generated MLE12 cells stably expressing either mLANA or kLANA (Fig 7D) and infected the cultures with a BAC16-derivative virus with a deletion encompassing the LANA-encoding region (KSHV-BAC16Δ73). Due to relatively low titers of the KSHV-BAC16Δ73 virus (∼1% initial infection efficiency) we were unable to perform ChIP-Seq analysis immediately after infection. However, as shown in the upper panels of Fig 7A, ChIP-qPCR performed after 5 days p.i. did not indicate any significant changes of H3K4-me3 or H3K27-me3 patterns across a number of human and viral marker loci. In particular, all viral loci that had acquired significant levels of H3K27-me3 in KSHV wt infected cells did so in transcomplemented KSHV-BAC16Δ73, regardless of whether cells were expressing kLANA or mLANA.

**Fig. 7:**
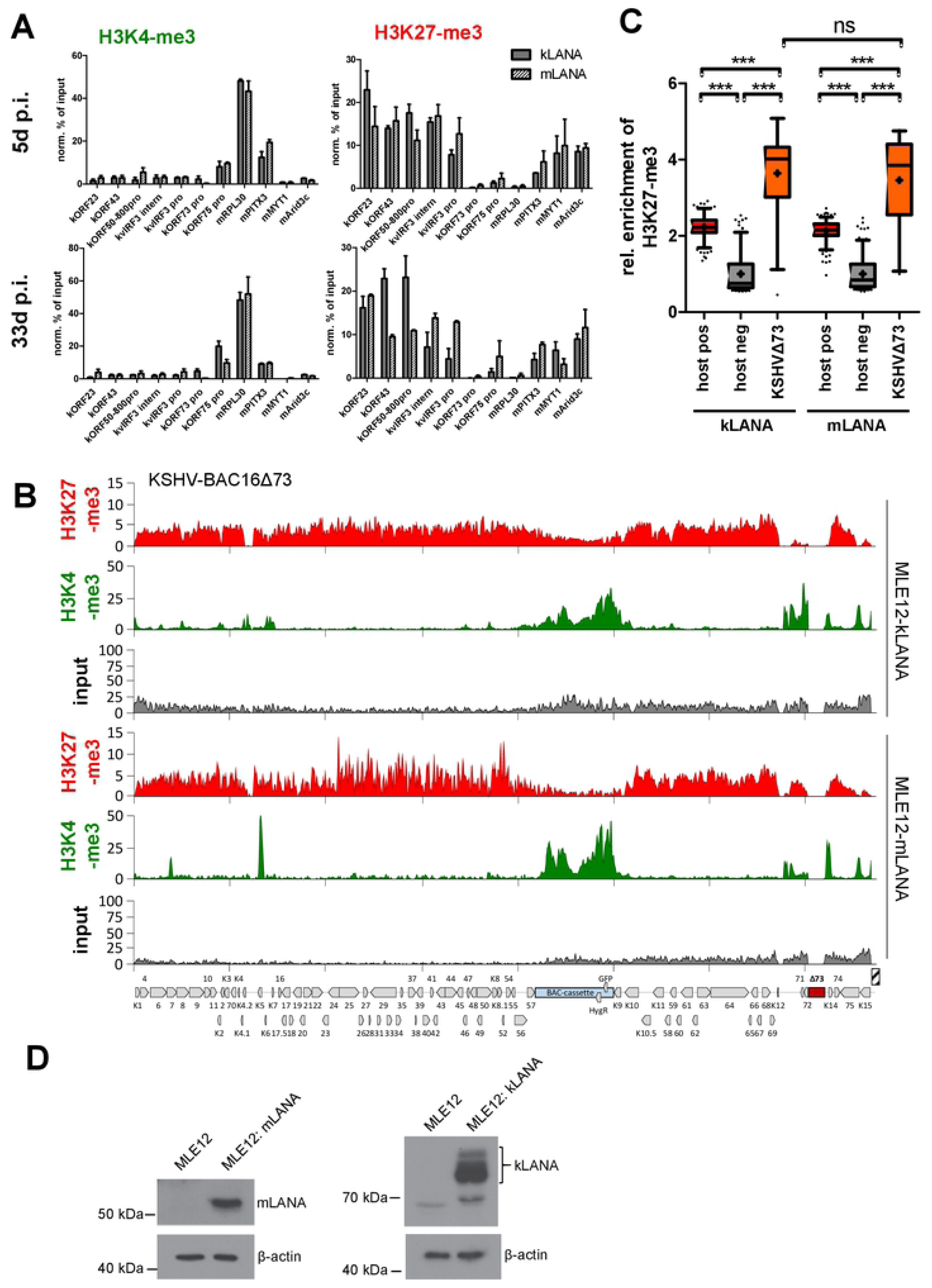
KSHV-BAC16Δ73 genomes trans-complemented by mLANA do not lose the ability to rapidly recruit H3K27-me3 marks. (A) ChIP-qPCR analysis of H3K4-me3 and H3K27-me3 in KSHV-BAC16Δ73-infected MLE12 cells that had been stably transduced with lentiviral mLANA or kLANA expression constructs. Cells were cultured in the presence of hygromycin to enrich for KSHV-BAC16Δ73-infected cells and ChIP was performed at day 5 (n>=2, top panel) or day 33 post infection (n>=2, bottom panel). Data are represented as mean ± SEM. (B) H3K27-me3 and H3K4-me3 ChIP-Seq analysis of material harvested after 33 days from KSHV-BAC16Δ73-infected kLANA (top panels) or mLANA (bottom panels)-expressing MLE12 cultures. Lower input coverage across the left half of the KSHV genome in MLE12-mLANA cells (see bottom panel) suggests loss of sub-genomic material from a fraction of episomes, similar to what has recently been observed with KSHV mutants expressing an oligomerization-deficient kLANA protein [25]. For visualization purposes ChIP-seq tracks were therefore normalized for using mean input coverage values. Non-normalized coverage data as used for statistical analysis is given in S1 Dataset. (C) Statistical analysis of H3K27-me3 enrichment on BAC16Δ73 episomes calculated as described in the legend to figure 4. Significance was calculated by 1way ANOVA testing (F = 431.2, df = 838). The difference between BAC16ΔLANA in kLANA vs. mLANA expressing cells was not significant (ns). (D) Western blot analysis of mLANA and kLANA expression in stably transduced MLE12 cells.

Under antibiotic selection, kLANA as well as mLANA-expressing cells (but not the parental ML12 cells) yielded hygromycin-resistant cultures, as was expected based on the previously reported ability of mLANA to support maintenance of episomes harboring KSHV terminal repeats [21]. We repeated our ChIP-qPCR analysis and additionally performed ChIP-Seq from the bulk-selected cultures at 33 days p.i., a similar late time point as in our previous MHV-68Δ50 or MHV-68Δ50-kLANA experiments. As shown in the lower panels of Fig 7A, ChIP-qPCR suggested that KSHV-BAC16Δ73 had maintained patterns of H3K4-me3- and H3K27-me3-positive loci as already observed after 5 days if infection. These results were furthermore confirmed by the ChIP-seq data presented in Fig 7B, which in both cases yielded very similar patterns of distinct H3K4-me3 peaks and broad, anti-correlated zones of H3K27-me3 as previously observed during KSHV wild type infection of MLE12 cells (Fig 5A; note that the additional peaks near the center in Fig 7B map to the bacmid cassette in BAC16 which constitutively expresses GFP and hygromycin resistance markers). Statistical analysis of H3K27-me3 levels on KSHV-BAC16Δ73 likewise indicated that KSHV-BAC16Δ73 episomes were able to efficiently attract H3K27-me3 in kLANA as well as mLANA-expressing cells (Fig 7C). The only appreciable difference was that input coverage across the left half of the KSHV genome was noticeably lower in MLE12: mLANA cells (see bottom panel in Fig 7C), an observation which is reminiscent of the loss of sub-genomic material recently observed with KSHV mutants expressing an oligomerization-deficient kLANA protein [25]. Nonetheless, our experiments consistently demonstrate that both KSHV and MHV-68 genomes maintain original dynamics of H3K27-me3 acquisition when (trans-)complemented with a non-syngeneic LANA protein, underlining our previous notion that cis-acting mechanisms are of pivotal importance and suggesting that kLANA does not possess unique properties which mediate rapid PRC recruitment.

### Analysis of MHV-68 infected splenocytes suggests that wildtype genomes in long-term latency pools are marked by H3K27-me3 *in vivo*

While our data show that MHV-68 genomes are unable to efficiently acquire H3K27me3 in a rapid fashion, the slight anti-correlation between H3K27-me3 and H3K4-me3 profiles in long-term infected cultures suggested that a small fraction of MHV-68 genomes nevertheless attract PRC, either as a result of an effective (but rare) stochastic event, or a constitutive (but generally weak) recruitment mechanism. If so, one may hypothesize that immune pressure *in vivo* could select or enrich for polycomb-repressed genomes during establishment of latency reservoirs. To investigate this possibility, we infected two mice with wildtype MHV-68 and isolated splenocytes and latently infected B-cells from the spleens 17 days after infection. Since the fraction of infected cells among the total population is very low we subjected the chromatin directly to ChIP-qPCR for H3K4-me3 and H3K27-me3 (Fig 8A), using a set of MHV-68 and host-specific primers from our previous *in vitro* infection experiments. Overall, the ChIP-qPCR results from both mice were highly concordant for both histone marks and across all loci, strongly arguing for the validity of the data. As shown in the top panel of Fig 8A, in both mice H3K4-me3 was readily detectable in three of the regions which had also tested positive in MHV-68Δ50 infected MLE12 cells (mORF72, M13 and downstream of M5), whereas the regions close to ORF4 and ORF64 were not enriched *in vivo*. Additionally, analysis of H3K27me3 demonstrate that *in vivo* latency reservoirs of MHV-68 indeed exhibit enrichment of repressive H3K27-me3 marks at most of the tested viral loci, at levels that were comparable to the endogenous positive control MYT1 (center panel of Fig 8A). The only loci which appeared completely devoid of H3K27-me3 were M13 and the region downstream of ORF72, i.e. the same loci which showed the strongest enrichment for H3K4-me3. The negative correlation of H3K4-me3 and H3K27-me3 at these loci indicates that the profiles originate from the same chromatin and do not represent differentially modified episomes.

**Fig. 8:**
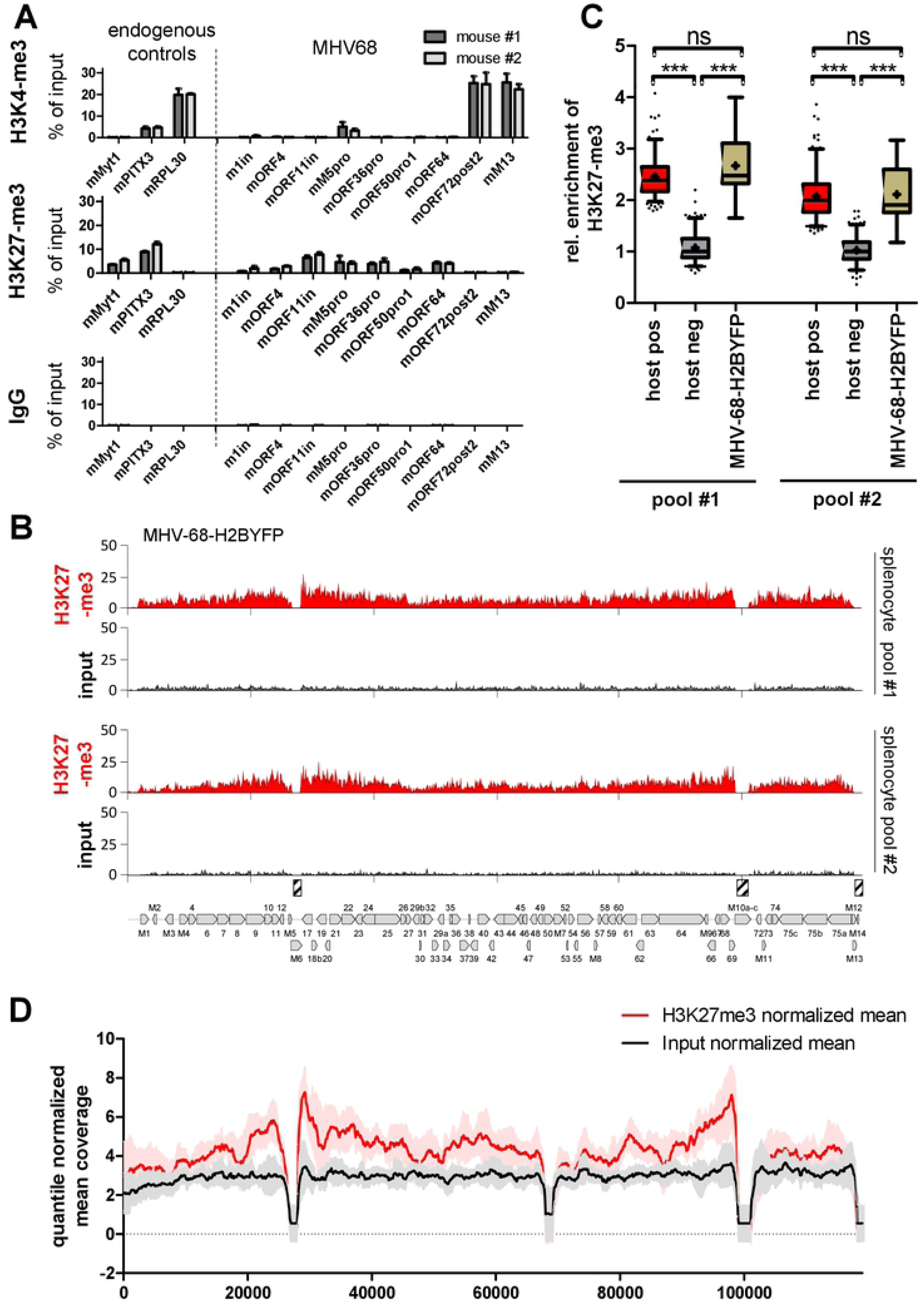
Histone-modification patterns of latent MHV-68 genomes *in vivo*. (A) ChIP-qPCR analysis of H3K27-me3 (top), H3K4-me3 (center) or IgG (negative control, bottom) in splenocytes harvest from two mice (#1 and #2) that had been intranasally infected with wildtype MHV-68 for 17 days, using MHV-68 or endogenous control primers as indicated. Data are represented as mean ± SEM of three independent ChIP replicates performed with the isolated chromatin from each mouse. (B) H3K27-me3 ChIP-seq coverage across the MHV-68 genome in two pools of splenocytes isolated from a total of six mice that had been intranasally infected with MHV-68-H2BYFP for 17 days. Splenocytes were FACS sorted for YFP expression prior to analysis. Due do the low number of positive cells, YFP-positive cells from three mice were pooled to generate pools #1 and #2. Approximately 5000 cells were subjected to low cell ChIP-seq using an H3K27-me3 specific antibody. Hashed boxes above the MHV-68 map indicate the position of repetitive regions (left and right internal repeat regions, as well as terminal repeat sequences) which had to be masked since they do not allow unique read mapping. (C) Relative enrichment of H3K27-me3 at viral episomes in splenocyte pools #1 and #2 was assessed using the same statistical method as described in the legend to Figure 4. (D) Normalized mean H3K27-m3 and input coverage from all MHV-68 ChIP-seq experiments performed in our study. Quantile normalized mean values and standard deviation (colored area) of H3K27-me3 and input tracks were generated from all MHV68-specific coverage tracks as given in S1 Dataset (ChIP MHV-68).

Encouraged by these results, we next aimed to enrich latently infected splenocytes for ChIP-seq analysis. For this purpose, we infected six mice with MHV-68-H2BYFP, a recombinant virus expressing an EYFP-H2B fusion gene for *in vivo* tracking of MHV68-infected cells [26], and performed FACS sorting of splenocytes isolated after 17 days of infection. As the recovered cell numbers per animal (approximately 3000) were too low for individual analysis we pooled cells from three animals each and performed RNA-seq as well as ultra-low input ChIP-seq for H3K27-me3 on the resulting pools. Transcriptomic Immgen cluster analysis of the 200 most highly expressed host genes in YFP-positive splenocytes confirmed that the cells showed typical expression patterns of germinal center B cells, as expected for authentic splenic latency reservoirs (S4 Fig). Mapping of RNA-seq reads to the MHV-68 genome furthermore revealed readily detectable transcription of ORF73 (the gene encoding mLANA) together with viral expression patterns that were highly restricted when compared to S11E or productively infected MLE12 cells (See S2 Fig D and E for coverage plots and expression analysis). Although gene expression profiles suggest the presence of a reactivated sub-fraction of cells, the splenocytes clearly cluster with MHV-68Δ50 infected MLE12 cells and away from S11E or lytically infected MLE12 cultures (S2 Fig E), indicating successful isolation of latency pools.

As shown in Fig 8B, analysis of H3K27-me3 in each of the two replicates indeed revealed global enrichment patterns that, in accord with our previous ChIP-qPCR analysis of wt-infected mice, reached similar levels as the endogeneous positive controls (Fig 8C). Again, the region encompassing M13 and (to a lesser degree) the regions downstream of ORF72 showed lower levels of H3K27-me3. Although the limited available material did not allow us to perform ChIP-seq for activation-associated marks, this observation suggests that episomes in long term latency reservoirs share at least some of the H3K4-m3 marks observed in our *in vitro* experiments.

Taken together, although the levels of H3K27-me3 enrichment did not reach those observed for latent KSHV episomes, the widespread distribution of H3K27me3 as well as its partial anti-correlation with H3K4-me3 is highly reminiscent of latent KSHV epigenome patterns. This suggests that polycomb-repression may indeed play a more concise role in *in vivo* latency reservoirs of MHV-68, potentially due to immune pressure that selects for cells harboring silenced episomes.

### Binding of KDM2B and acquisition of H2AK119-ub during early infection suggest that KSHV attracts polycomb repressive complexes via the non-canonical recruitment pathway

Our previous results suggested that KSHV rapidly attracts H3K27-me3, whereas MHV-68 may do so in a manner that proceeds stochastically or with low efficiency, and that cis-acting features underlie the observed differences. We therefore sought to identify sequence features that could potentially explain our observations. Regarding potential recruitment mechanisms in MHV-68, when overlaying normalized H3K27-me3 patterns from all our MHV-68 experiments (Fig 8D) we noticed that the regions flanking the left and right internal repeats (IR1 and −2 in the following) consistently reached the highest enrichment levels. This suggests that PRCs may initially be recruited to the internal repeats (note that, due to their repetitive nature, ChIP-seq reads cannot be mapped to the IR elements themselves) and subsequently spreads into adjacent regions, with the region to the right of IR2 being more refractory to spreading due to the presence of H3K4-me3 marks. In contrast, our previous temporal analysis [8] suggests rapid global acquisition of H3K27-me3 methylation to KSHV genomes, rather than spreading from nucleation sites. When we searched for similarities between the KSHV and the MHV-68 IR elements that are not shared by the remainder of the MHV-68 genome, the prime characteristics that emerged were those of CpG islands. As shown in S5 Fig B, the MHV-68 genome is highly CpG suppressed (suppression index 0.43) and exhibits a general GC content of 47%. Only relatively short regions meet the criteria of CpG Islands, chief among them the two IR regions. In contrast, nearly the entire KSHV genome (average GC content 54%, CpG suppression index 0.84) registers as a single, continuous CpG island (S5 Fig A). This observation was of particular interest since CpG islands have emerged as the prime PRC recruitment signal in vertebrate cells during the recent years [reviewed in 27, 28]. In this pathway, the non-canonical PRC1.1 complex is directly recruited via binding of its subunit KDM2B (a H3K36-specific demthylase [29]) to unmethylated CpGs, followed by ubiquitination of H2A lysine 119 (H2AK119-ub) [30-33]. Given the fact that herpesviruses package epigenetically naive DNA, and that KSHV episomes do not acquire substantial levels of DNA methylation until several weeks post infection [6], we therefore tested the possibility that de novo-infecting KSHV genomes recruit KDM2B and performed ChIP-seq analysis at 24h post infection. As shown in Fig 9A, we indeed detected abundant and global binding of KDM2B to KSHV episomes, at levels that were comparable to those seen on the most significantly enriched positive host loci (Fig 9B). Investigation of the signals across the host genome furthermore confirmed the expected binding patterns at CpG Islands [30].

**Fig. 9:**
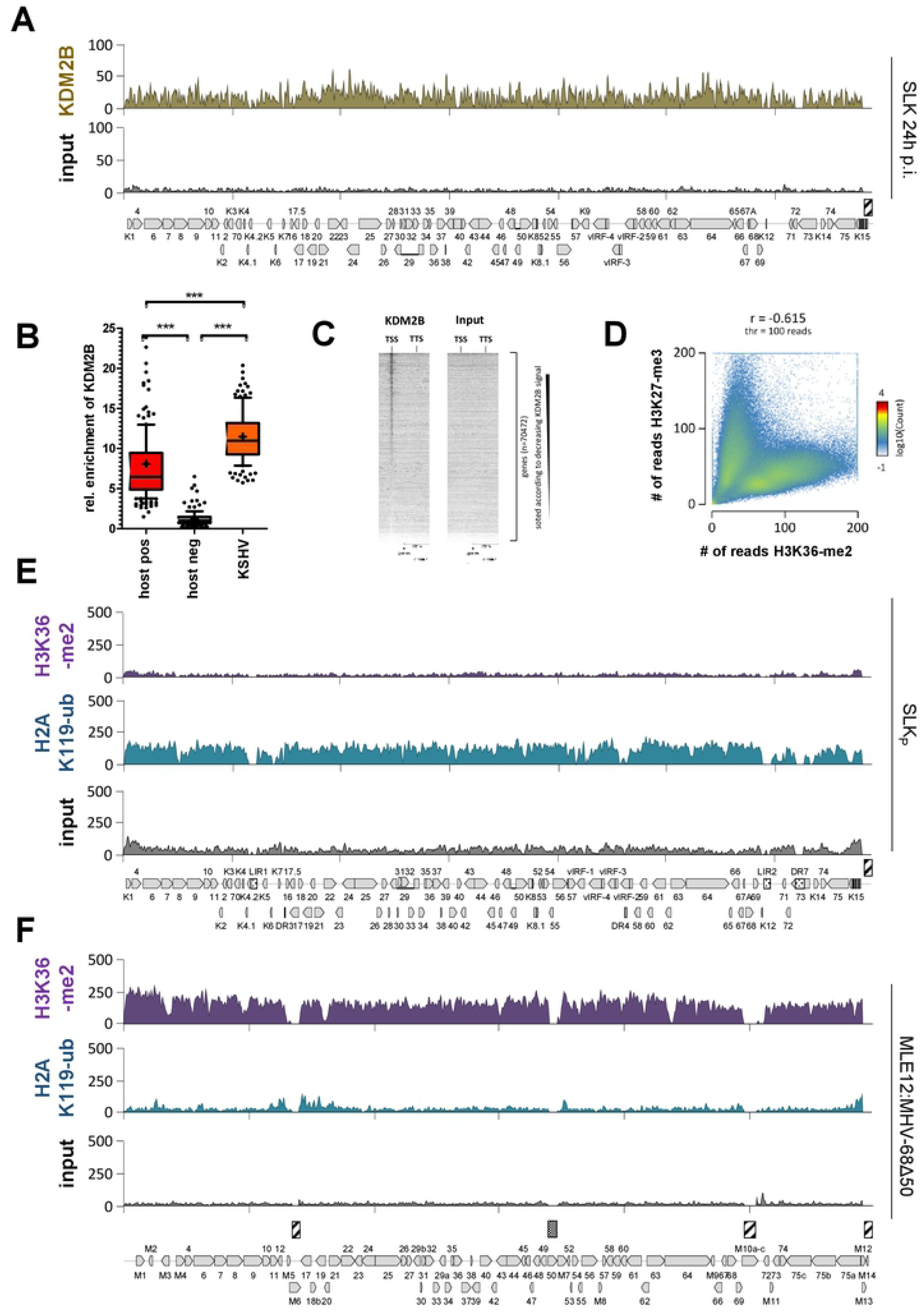
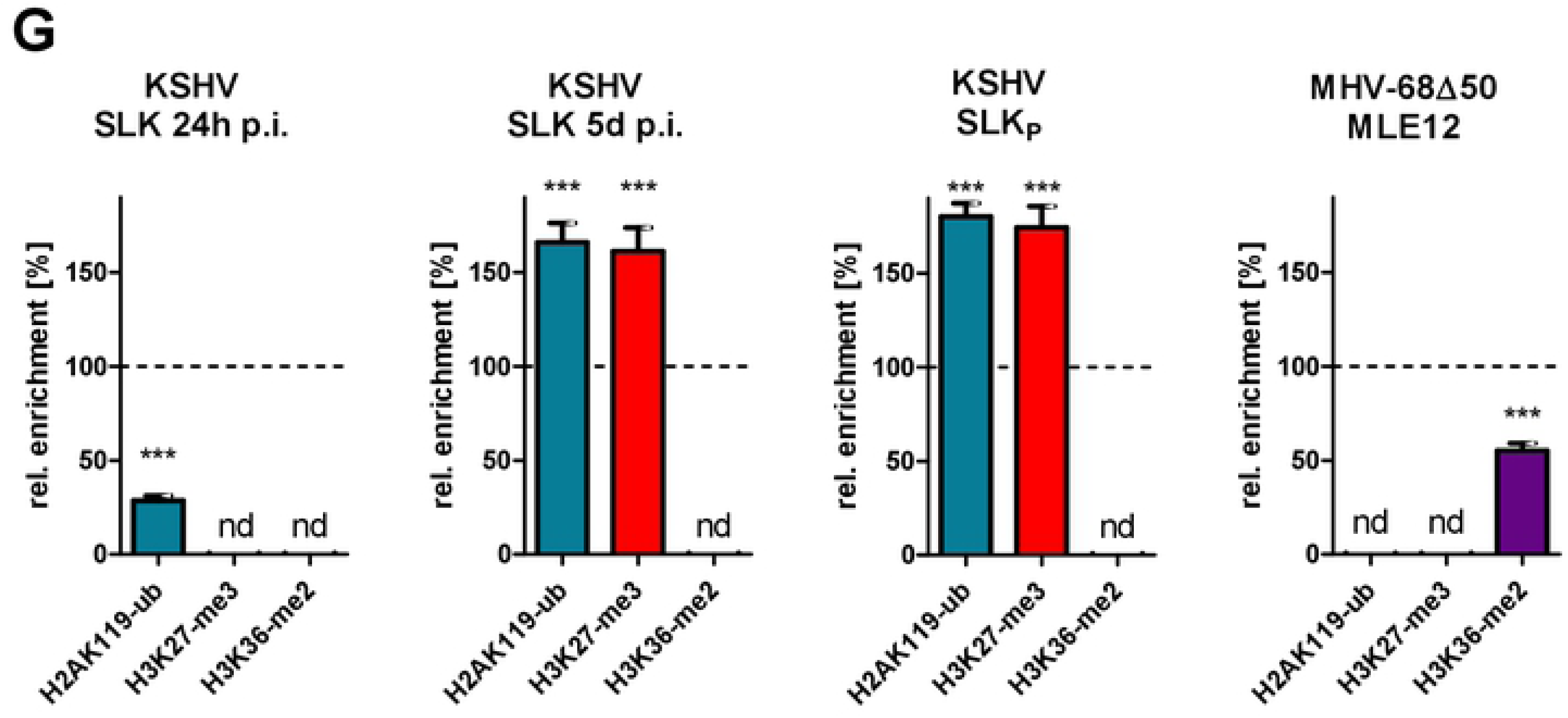
KDM2B recruitment and acquisition of PRC1-associated histone modifications by KSHV and MHV-68Δ50 genomes. (A) KDM2B ChIP-seq coverage (top) or input (bottom) profiles across the KSHV genome in SLK cells at 24 hours post-infection. (B) Relative enrichment of KDM2B on *de novo* infecting KSHV episomes measured at 24 hours post infection. Enrichment was quantified similar to method described in the legend to Figure 4. KDM2B enriched human positive control regions were detected by MACS peak calling. Significance was calculated by 1way ANOVA testing (F = 372.5, df = 508). (C) Heatmap of KDM2B enrichment at transcriptional start sites (TSS) of human genes generated from KDM2B ChIP-seq data at 24 hours post infection with KSHV infection. The Y-axis represents 70,474 individual length normalized annotated transcripts (from TSS to TTS) of human genes, and the X-axis represents the 200 % surrounding of the length-normalized genes. Data was sorted according to decreasing KSM2B signal at the TSS. (D) Global anti-correlation of cellular H3K27-me3 and H3K36-me2 patterns across the human genome. The 2D histogram shows the number of genomic 10000 bp windows with the combination of the number of reads described on the axes in SLKp cells. The colored bar to the right side illustrates the number of windows with each combination of reads in the two datasets. (E+F) ChIP-seq coverage for H3K36-me2 (top) and H2AK119-ub (center) or input (bottom in each panel) from (E) stably KSHV-infected SLK_P_ cells or (F) long-term MHV-68Δ50-infected MLE12 cells. (G) Analysis of significant enrichment of H2AK119ub, H3K27me3 and H3K36me3 on KSHV genomes in (left panel) SLK cells after 24 hours of infection, (2nd panel from left) SLK cells after 5 days of infection or (2nd panel from right) SLKp cells, or (right panel) MHV68 genomes in long-term MHV68-Δ50-infected MLE12 cells. Based on the statistical method described in the legend to Figure 4, normalized enrichment was calculated relative to the median value of the host loci that showed strongest and weakest enrichment (set to 100% and 0%, respectively) for each modification. Significance indicators (asterisks) are shown for those modifications for which median enrichment along the viral genome was significantly above levels observed for the cellular background. ‘nd’ (not detectable) demarks modifications in which enrichment was not significantly different from (or signficantly lower) than in the negative control regions. Data from MLE12 and SLKp cells correspond to those shown in E and F above, or (for H3K27-me3) those in Fig 4. Coverage tracks for SLK cells at 24 h.p.i and 5 d.p.i. are provided in S6 Fig. Non-normalized statistical data for each of the modifications and cell lines shown in this figure are provided in S7 Fig.

Unfortunately, we were unable to perform similar experiments with MHV-68 infected cells since later lot numbers of the KDM2B antibody did not show expected enrichment patterns on the host genome, rendering the fact that we did not detect any signals on the MHV-68 genome with these antibodies inconclusive. In the absence of a working ChIP-grade KDM2B antibody, we therefore investigated patterns of H3K36-me2, a histone mark that is the direct target of KDM2B’s demethylase activity. Given the observed binding to KSHV genomes, we expected that KSHV episomes should exhibit significantly lower levels of H3K36-me2 compared to MHV-68. As shown in the top panels of Fig 9E (KSHV) and F (MHV-68), this is indeed the case: Whereas KSHV exhibits very low signals of this histone mark in SLKp cells, levels on MHV-68 episomes in MLE12:MHV-68Δ50 cells were significantly higher and reach levels that were comparable to those of endogenous positive control loci (see statistical analysis in S7 Fig). Global analysis of H3K36-me2 methylation patterns across the host genome (Fig 9D) furthermore verified general anti-correlation with H3K27-me3 [34], thus confirming the validity of our data.

We next investigated global patterns of H2AK119-ub, the ubiquitination mark deposited by PRC1 complexes. As shown in the center panel Fig 9F, although a slight enrichment near the CpG-rich left internal repeat unit was noticeable, MHV-68Δ50 genomes overall showed H2A-K119ub levels that were not significantly above those of the host background (S7 Fig). In contrast, as expected KSHV genomes in SLKp cells were highly enriched for this mark (Fig 9E, center panel). If KSHV genomes acquire H3K27-me3 marks via KDM2B and the non-canonical PRC2 recruitment pathway, then global acquisition of H2AK119-ub would be expected to occur simultaneously with, or even prior to H3K27-me3 accumulation, and should also preclude H3K36-me2 acquisition during early infection. Indeed, a previous report by Toth and colleagues found that binding of RYBP, a component of non-canonical PRC1.1 complexes, precedes that of the PRC2 component EZH2 at a number of investigated loci [7]. To directly compare *de novo* acquisition of PRC1- and PRC2-dependent marks on a global level, we performed ChIP-seq for H2AK119-ub, H3K27-me3 and H3K36-me2 in SLK cells after 24 hours and 5 days post of infection. The graphs in Fig 9G show global enrichment of each mark across the viral genome relative to average levels of the 200 most strongly enriched host loci (set to 100%; see S6 and S7 Figs for full coverage plots and results of the statistical analysis). Statistical analysis of data from SLKp and MHV-68Δ50-infected MLE12 cells are shown to the right for comparison. In accord with our previous observation that H3K27-me3 marks require 48-72h to accumulate [8], we observe that average H3K27-me3 levels are very high in SLK cells after 5 d.p.i. or in long-term infected SLKp cells, but that enrichment over cellular background is not yet significant at 24h p.i. (three left plots in Fig 9G). In contrast, global enrichment of H2AK119-ub indeed is already highly significant after 24h p.i., and further increases through the 5d p.i time point. As expected, H3K36-me2 enrichment on KSHV episomes was not observed at any time point, whereas this mark was substantially enriched on MHV-68Δ50 genomes in MLE12 cells (right plot in Fig 9G). Taken together, these data strongly support the hypothesis that KSHV genomes recruit PRC via the non-canonical pathway, likely as a result of their high density of unmethylated CpGs.

## Discussion

The factors determining the epigenetic fate of nuclear viral DNA remain poorly understood. Herpesviruses represent attractive model systems to study such processes, given that they package unmethylated and nucleosome-free genomes and thus must newly establish a suitable chromatin landscape upon each round of infection to support either their productive phase or latency. To address the role of polycomb group proteins in this process, we performed a comparative analysis of epigenetic modifications acquired by latent KSHV and MHV-68 genomes. Our comparison involves two viruses which share many evolutionary conserved traits on the genomic and protein level, yet, at least in vitro, show distinct biological behavior after *de novo* infection: whereas KSHV establishes latency in a wide variety of cell types and lines, MHV-68 infected cells are much more prone to undergoing lytic replication. We hypothesized that these differences may be reflected on the epigenetic level and that, if so, our comparative analysis may open new avenues towards elucidating the viral features, which regulate PRC recruitment. Indeed, our initial analysis of S11E cells suggested that MHV-68 attract little, if any, H3K27-me3 marks. Even though these cultures harbored a significant fraction of lytic cells, the fact that H3K27-me3 profiles on latent KSHV genomes remain readily detectable even in lytically-induced BCBL1 cultures [11] suggests that MHV-68 genomes in the latent fraction of cells do not carry abundant H3K27-me3 marks. However, as presence of lytic cells nevertheless complicates interpretation of these observations, we proceeded to investigate the epigenetic landscape of *de novo* infecting MHV-68 genomes that are deficient for the master lytic transactivator encoded by ORF50. At first, it may seem counterintuitive to employ a state of forced latency to investigate the factors which may favor latency in the first place. However, these factors are likely to act prior to lytic cycle entry and ORF50 expression. In support of this, Toth and colleagues recently reported that an ORF50/Rta deficient KSHV attracts H3K27-me3 marks indistinguishably from its wildtype counterpart [13], an observation which is in line with our own unpublished results. Hence, there is every reason to believe that the MHV-68Δ50 mutant provides an appropriate measure of the efficiency with which latently replicating episomes recruit PRCs. Indeed, our results demonstrate that MHV-68Δ50 exhibits a latent expression profile, but nevertheless does not attract abundant H3K27-me3 marks. Importantly, we show that this is independent of the host cellular background, as co-infecting KSHV episomes rapidly acquire H3K27-me3 marks at levels that are on par or above the most significantly enriched host loci.

Interestingly, although H3K27-me3 levels on MHV-68Δ50 were low, we detected anti-correlation with activation marks, suggesting that a subset of episomes may be targeted by PRC2. Considering this, our finding that H3K27-me3 was acquired to a higher extent in latently infected splenocytes isolated from mice at day 17 post infection indicates that PRC recruitment to MHV-68 episomes is possible, but may result from an inefficient or stochastic process. This suggests a model in which MHV-68 may have evolved to undergo rapid expansion via lytic replication in the lung of infected animals, followed by selection of cells harboring polycomb-repressed episomes in the spleen. Since we detected a low-level transcriptional signature of lytic genes in MHV-68Δ50-infected MLE12 cells, we postulate that stochastic firing of lytic promoters may generate immunogenic pressure for the selection of latently infected cells with tighter repression patterns. Of course, it is also possible that differences in the cellular background of splenocytes may allow MHV-68 to acquire H3K27-me3 more efficiently. If so, however, the fact that we did not observe any H3K27-me3 marks in S11E cells (a B cell line) argues against the hypothesis that it is the epithelial background of MLE12 cells *per se* which prevents acquisition of H3K27-me3. More importantly, we find that co-infecting KSHV episomes efficiently, rapidly and globally attract H3K27-me3 marks in MLE12 cells. Hence, the mechanisms that allow rapid PRC recruitment to KSHV genomes must be fully functional in MLE12 cells, yet do not act upon MHV-68 genomes. This allows us to draw important conclusions with regard to the nature of pathways attracting PRCs to KSHV. However, owing to the limitations of available MHV-68 latency models, it does not permit us to exclude the possibility that MHV-68 may employ different (and potentially splenocyte-specific) mechanisms in authentic *in vivo* latency reservoirs. Given that CpG-dependent pathways have emerged as a fundamental recruitment mechanism in mammals, we propose that the initial recruitment to viral genomes generally proceeds via the non-canonical pathway, but that KSHV’s high CpG frequency serves to ensure rapid silencing in a manner that is largely independent of the cellular background, whereas MHV-68 may additionally require, for example, an environment that allows more efficient spreading of PRC from initial seed sites. Certainly, further studies will be required to elucidate the mechanism that may govern PRC-dependent repression of MHV-68 *in vivo*.

Although MHV-68 and KSHV episomes behave differently regarding PRC recruitment, they both acquire a number of distinct activation marks. Upon in vitro infection, activating histone modifications were acquired within the first days and most (but not all) of the peaks persisted throughout long-term infection, a feature which is reminiscent of observations made previously in the KSHV system [6, 8]. Our transcriptional analysis (S3 Fig, S5 Dataset) suggests that, as in KSHV [11], the activation-marked regions are preferentially associated with immediate early gene promoters. The fact that most activation marks are not associated with abundant transcription suggests that additional viral or host factors (including, but not necessarily limited to, ORF50/Rta) are required to permit full activation. We therefore suspect that the role of persistent activation marks is to keep immediate early promoters in an open chromatin configuration during latency, such that the virus can rapidly respond when such factors become available. Indeed, in methylated DNA immunoprecipitation experiments (MeDIP-seq, see S2 Protocol) we have observed that, while the MHV68 genome is principally subject to DNA methylation in long-term infected MLE12 cells (including, as previously reported [17, 18], at the distal ORF50 promoter), loci harboring persistent H3K4-me3 peaks remain negative. Whether such marks indeed facilitate reactivation from latency, however, remains to be established.

Finally, what are the features which mediate rapid recruitment of PRCs to KSHV episomes, and why does MHV-68 (at least in our model systems) exhibit such fundamentally different behavior? While further experiments will be required to work out the precise details, our study provides a framework which limits the possible options and suggests a potential mechanism. Firstly, our co-infection experiments clearly demonstrate that it is not a lack of specific host factors which preclude rapid PRC recruitment to MHV-68 genomes. At the same time, these data also show that latently expressed, trans-acting viral gene products encoded by MHV-68 do not interfere with efficient acquisition of H3K27-me3 by KSHV. Conversely, we can also conclude that expression of KSHV products is not sufficient to convey rapid PRC recruitment to MHV-68 episomes. We therefore considered two, mutually non-exclusive options: that polycomb recruitment may be regulated by viral cis-acting gene products that only act upon their parental episomes, or by unique genomic features that are present in the KSHV, but not the MHV-68 genome. A prime candidate for the former was the KSHV-encoded kLANA, given that a recent study [13] reported that a LANA-deficient virus fails to acquire H3K27-me3 marks. The study furthermore found kLANA to decorate the entire viral episome (likely via its unspecific chromatin binding activity) and to form high molecular-weight complexes with PRC2 components. Although it is presently unclear whether this complex formation reflects a direct interaction with PRC2 components or is a consequence of kLANA’s chromatin binding ability per se, this suggests a model in which LANA may directly promote PRC recruitment.

Our experiments with recombinant viruses show that substitution of mLANA by kLANA does not convey the ability to rapidly acquire H3K27-me3 marks to MHV-68 genomes. Reciprocally, KSHV episomes that are maintained by mLANA instead of kLANA do not lose this ability. Hence, whatever function of kLANA is required for H3K27-me3 acquisition must be shared with mLANA, but at the same time must be insufficient to mediate rapid PRC recruitment in the context of the MHV-68 genome. This suggests one of the core features of the LANA proteins, such as the ability to support licensed DNA replication, as a pre-requisite for H3K27-me3 acquisition, a hypothesis which we are currently investigating. We would like to point out that such models do not exclude the possibility that interaction of kLANA with PRC2 components may additionally facilitate H3K27-me3 acquisition. However, they suggest that the primary trigger of rapid PRC recruitment is distinct [13]. The fact that LANA has to accumulate in newly infected cells before it can bind to viral chromatin also suggests that other factors are critically involved in the recruitment process.

Based on our results, we propose that these factors involve the same genomic sequence features which also mediate PRC recruitment to host loci [reviewed in 27, 28]. Traditionally, polycomb repression has been thought to result from a strictly sequential process: first, binding of PRC2 and EZH2-dependent tri-methylation of H3K27, and second, recognition of H3K27-me3 marks by the PRC1 component Cbx and subsequent H2AK119 ubiquitination and transcriptional repression. Certain transcription factors such as YY1 can recruit PRC2 in mammals, but the fact that KSHV genomes acquire H3K27-me3 marks in a gradual and uniform manner argues for a global recruitment process, rather than spreading from discrete transcription factor binding sites [8]. Interestingly, the traditional model is increasingly being challenged by studies suggesting that PRC2 binding and H3K27 methylation can also represent a secondary event that is preceded by direct PRC1 recruitment [reviewed in 27, 28, 35]. The prime genomic feature associated with PRC1 recruitment is high density of non-methylated CpG dinucleotides, and CpG density and methylation states are in fact good predictors of PRC2 recruitment in mammals [36]. Accumulating evidence suggests that the H3K36 demethylase KDM2B, a component of the non-canonical PRC1.1 complex which can bind to un-methylated CpG motifs via its CXXC zinc finger domain, is a major factor directing polycomb complexes to CpG islands [30-33]. A recent study furthermore demonstrated that gene silencing triggers efficient recruitment of PRC2 to CpG islands in a genome wide manner, suggesting that PRC2 by default binds to non-transcribed CpG islands [37].

Considering the above, it is intriguing that the prime genomic feature which distinguishes KSHV from MHV-68 is CpG suppression, with MHV-68 showing much more severe suppression compared to KSHV (suppression index of 0.43 and 0.82, respectively). Indeed, when using common definition criteria almost the entire KSHV genome scores as a single, contiguous CpG island, whereas only short interspersed segments meet the requirements in the MHV-68 genome (S5 Fig). The fact that KDM2B efficiently binds to de novo infecting KSHV genomes, and that H3K36-me2 levels are substantially lower on KSHV compared to MHV-68 genomes further supports the hypothesis that CpG frequency may drive rapid accumulation of H3K27-me3 marks on KSHV episomes via the non-canoncial PRC1.1. Certainly, additional studies will be required to investigate the role of KDM2B and non-canonical PRC complexes in latency establishment. For example, thus far we have been unable to generate viable knockout or knockdown cells in which KDM2B levels were substantially reduced, and the overall impact of KDM2B and PRC1.1 activity on H3K27-me3 accumulation and silencing of lytic KSHV genes therefore remains unknown. We are currently targeting other components of PRC1.1 to further investigate this issue.

In the meantime, the working model presented in Fig 10 may provide a potential explanation of the observations made in the present study, and may also serve as a basis to inform future studies by our and other groups. We propose that, for KSHV as well as MHV-68, one of the earliest events after nuclear entry is the binding of sequence-specific transcription factors of hitherto unknown nature, leading to recruitment of chromatin modifiers (including H3K4 specific methyltransferases such as MLL and SET2) that demark the observed patterns of activation-associated histone modifications on latent genomes. In KSHV, the high density of unmethylated CpG motifs results in rapid acquisition of non-canonical PRC1.1 via its KDM2B component and subsequent PRC2 recruitment. Secondary recruitment of PRC2 has recently been shown to be promoted by H2AK119-ub marks deposited by PRC1 [33], and a similar order of events for KSHV genomes is suggested by the data presented in Fig 9 together with the previous finding that binding of RYBP, a component of non-canonical PRC1 complexes, precedes that of the PRC2 component EZH2 [7]. However, given recent reports of methylation sensitive DNA binding by PRC2-accessory proteins of the Polycomblike family [38-40], PRC2 may also be able to directly bind to non-methylated KSHV DNA.

**Fig. 10:**
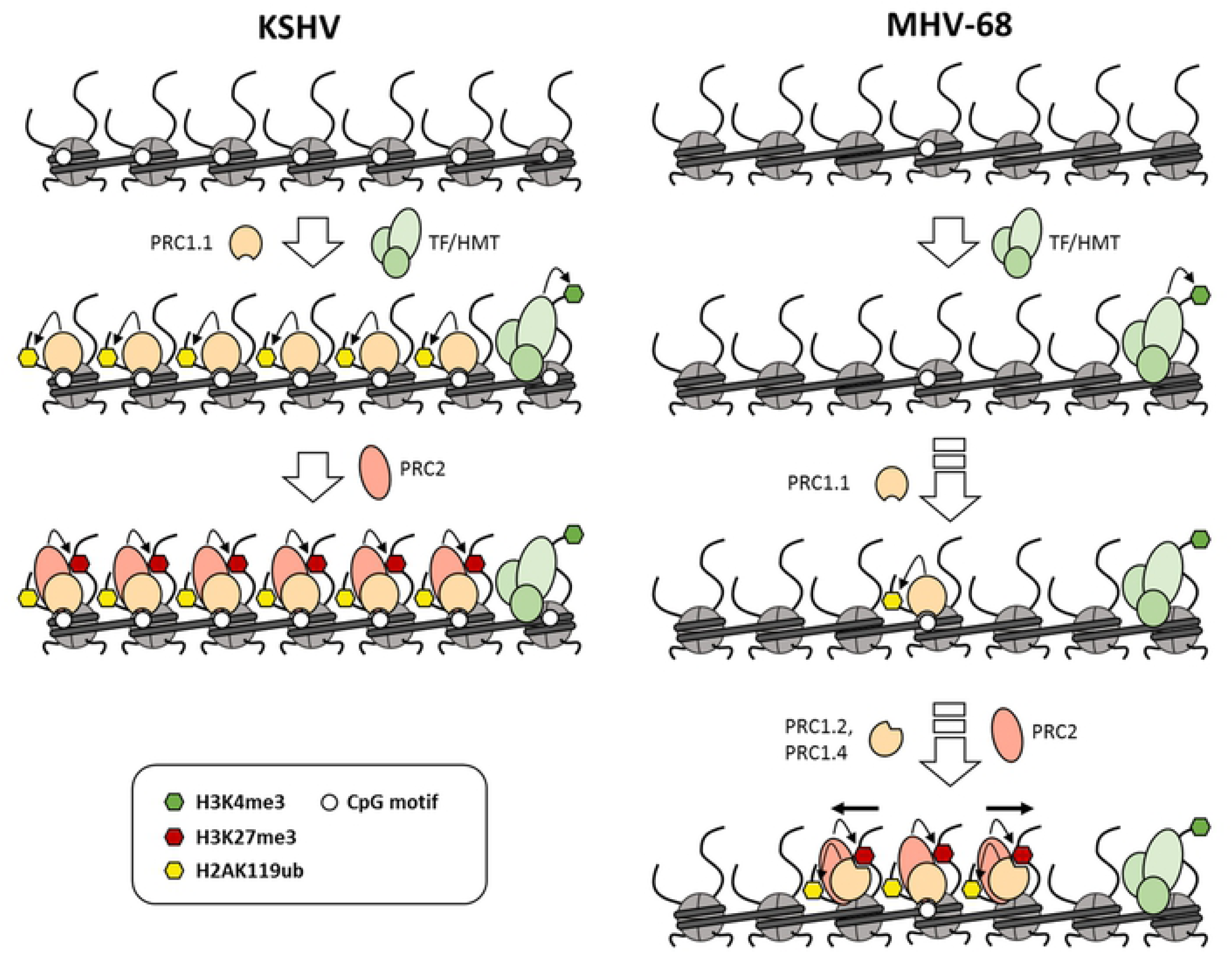
Model of PRC1 and −2 recruitment to KSHV and MHV-68 genomes. Immediately after nuclear entry, sequence-specific transcription factors bind to KSHV as well as MHV-68 genomes and lead to deposition of activation-associated H3K4-me3 marks. In KSHV (left panel), high density of unmethylated CpG motifs mediates rapid acquisition of the non-canonical PRC1.1 complex, followed by PRC2 recruitment as a secondary event (alternatively, PRC2 might also be directly recruited to CpG-rich DNA). Owing to the lower CpG frequency of MHV-68 genomes (right panel), only relatively short sequence segments which exhibit characteristics of CpG islands (such as the internal repeat regions) are initially acquire PRC1.1 complexes in a delayed (depicted) or stochastic manner. Once established, canonical PRC occupancy and associated histone modifications may slowly spread from these sites.

While the lower CpG frequency of MHV-68 genomes does not allow rapid genome-wide recruitment of PRC complexes, our data suggest that the internal repeat regions may nevertheless serve as seed regions for polycomb acquisition. This hypothesis is also supported by the fact that the left internal repeat region has been previously found to be important for latency amplification in vivo [41]. In the right panel of Fig 10 we have therefore depicted a scenario in which PRC1.1 binds initially at CpG-rich (repeat) regions of the MHV-68 genome in a delayed manner (or as the result of a stochastically rare event), followed by spreading of PRCs and associated histone modifications sites via canonical complexes. As discussed above, whether or not PRC acquisition by MHV-68 is facilitated by splenocyte-specific factors remains to be established.

Given our results, we propose that PRC recruitment may represent a default response to nuclear, epigenetically naive viral genomes that harbor a high density of non-methylated CpG motifs, and that different viruses may have adopted different strategies to exploit, delay or evade this process. In this context, KSHV LANA may have evolved to further stimulate PRC2 binding and thus promote rapid latency establishment, whereas CpG depletion in MHV-68 genomes may reflect a lifestyle that is more prone to initial lytic replication in the lung, followed by selection of repressed genomes in splenic long-term latency reservoirs. Interestingly, KDM2B was recently also shown to bind to lytic EBV promoters and repress transcription [42], suggesting that EBV may also employ polycomb repression to support latency. Notably, however, the above does not mean that all CpG-rich DNA viruses will undergo polycomb repression, as they may have evolved other mechanisms to actively interfere with PRC recruitment. While it thus should be of great interest to study polycomb repression in other viruses, we expect that the genetically tractable system and evolutionary informed system presented here will be particularly useful in helping to decipher the molecular mechanisms which direct PRC to epigenetically naive, nuclear-invading DNA.

## Materials and Methods

### Cell lines and culture conditions

MLE-12 cells (ATCC; CRL-2110) and SLK_P_ cells (a pool of long-term infected single cell clones derived from SLK cells infected with the BCBL1 KSHV strain) [4] were grown in DMEM High Glucose (Gibco, Darmstadt, Germany) supplemented with 10% FCS, 2 mM L-Glutamine, 100 U/ml Penicillin and 100 μg/ml Streptomycin. S11E cells [16] and BCBL1 cells [43] were cultured in RPMI 1640 medium containing 10% FCS, 100 U/ml Penicillin and 100 μg/ml Streptomycin. SLK_P_ cells are a permanently KSHV positive cell line. The parental cell line SLK has been recently discovered to be a misidentified cell line (Cellosaurus AC: CVCL_9569) and was identified as a contaminating cell line (Caki-1; Cellosaurus AC: CVCL_0234) and is listed in *ICLAC.* Nevertheless, these cells have been shown to efficiently support KSHV latency as stated in several publications throughout the last years and therefore represent a suitable model cell line to study chromatinization of latent KSHV genomes. All cell lines were regularly tested for mycoplasma contaminations using LookOut® Mycoplasma PCR Detection Kit (Sigma-Aldrich, MP0035-1KT). Additionally ChIP-seq reads from input samples were aligned to mycoplasma reference sequences to exclude contamination.

### Generation of kLANA/mLANA expressing MLE12 cells

To generate lentiviral expression constructs for kLANA and mLANA, the respective ORFs were PCR amplified from KSHV and from MHV-68. PCR products were cloned into LeGOiC2 vectors [44]. Constructs were verified by Sanger sequencing. Lentiviral particles of LeGo-iC2-kLANA and LeGo-iC2-mLANA were produced in LentiX293 cells. These lentiviral particles were used to transduce MLE12 cells. During transduction 8 µg/ml polybrene was added. 8h post transduction medium was changes to complete DMEM. Cells were sorted multiple times to achieve a homogenous population of kLANA and mLANA expressing cells. Successful expression of kLANA and mLANA was verified by Western blot (see Fig 7D).

### Rodents

Female Balb/c mice (6-8 weeks old) were purchased from Charles River Laboratories (Sulzfeld, Germany) and housed in individually ventilated cages (IVC) during the MHV-68 infection period. All animal experiments were in compliance with the German Animal Welfare Act (German Federal Law §8 Abs. 1 TierSchG), and the protocol was approved by the local Animal Care and Use Committee (District Government of Upper Bavaria; permit number 124/08).

### Viruses and infection

#### MHV-68

Wildtype MHV-68 (ATCC; VR-1465) and MHV-68-H2BYFP [26] was propagated in MLE-12 cells by incubation with infectious virus supernatant resulting in a rapid productive lytic infection. Supernatant was harvested and filtered 72 to 96 hours post infection, when cells showed a visible cytopathic effect (CPE). Infection was performed by incubating 5×10^5^ MLE-12cells with 20 µl of prepared virus supernatant in 600 µl medium containing 8 µg/ml polybrene in a 6-well dish. Cells were cultured in EBM-2 medium without supplements (LONZA) for two hours followed by substitution with culture medium. For in vivo experiments, mice were inoculated intranasally (i.n.) with 5 x 10^4^ PFU of wildtype MHV-68 or MHV-68-H2BYFP. Prior to i.n. inoculation, mice were anesthetized with ketamine and xylazine. After 17 days, when latency is usually established in the majority of infected B-cells, mice were sacrificed and single splenocyte suspensions were prepared by forcing the prepared spleens trough a cell strainer into PBS. Cells were treated with RBC lysis buffer (Biolegend: #240301) and either directly fixed for ChIP or stained for dead cells using the Zombi NIR Fixable Viability Kit (Biolegend: #423106) as per the manufacturer’s instructions for subsequent FACS sorting (MHV-68-H2BYFP). Sorted cells were subjected to low cell ChIP and low cell RNA-seq analysis.

An MHV-68 ORF50 deletion mutant (MHV-68Δ50) was generated by ET-cloning of a GFP expressing MHV-68 BAC construct as described previously [45]. For this purpose, ORF50 was first partially replaced with a tetracycline (Tet) resistance gene flanked by FRT sites. Subsequently, the Tet resistance cassette was removed by FLP-mediated recombination, resulting in a deletion of nucleotides 67,957 to 69,326. The BAC-cloned genome was analyzed by restriction enzyme analysis with several restriction enzymes and high-throughput Illumina sequencing.

MHV-68ΔORF50 mutant virus was reconstituted on trans-complementing BHK-21 cells (ATCC; CCL-10) stably expressing ORF50. The latter were generated by transfection of BHK-21 cells with the eukaryotic expression vector pCR3 (Invitrogen) containing an EcoRI - AvaI fragment of MHV-68 (nucleotide positions 65,707 to 69,550). After transfection, cells were cultured in the presence of 500 µg/ml G418 (PAA). For virus reconstitution, the ORF50 trans-complementing BHK-21 cells were transfected with 2 μg of ΔORF50 BAC DNA using FuGENE HD Transfection Reagent (Roche, Mannheim, Germany). When cells showed a visible CPE, the supernatant was harvested and stored. Virus stocks were grown and titrated as described previously [45], except that ΔORF50 mutant virus stock was grown and titrated on the ORF50 trans-complementing BHK-21 cells. Infection was performed as described for MHV-68 (wt), however, we used a higher volume of viral supernatants (20 to 250 µl) since the mutant generally grew to lower titers compared to wild type. To investigate long-term infected cultures, MLE-12 cells were infected with trans-complemented MHV-68Δ50 virus expressing GFP from the BAC cassette [45]. Infected GFP-expressing cells were sorted approximately once per week using a FACS-Aria (BD) to maintain an almost 100% positive cell population for subsequent experiments. Loss rates were estimated to be approximately 5-10% per cell generation, as judged from the ratio of green versus non-fluorescent cells and the number of cell doublings (determined by counting cells during the sub-culturing). Note that these loss rates are only estimates as we did not perform FACS analysis at fixed intervals, or determined the fraction of virally infected cells in which the that GFP reporter may have be transcriptionally silenced.

To generate MHV-68Δ50-kLANA, the same strategy as decribed above for the MHV-68ΔORF50 mutant was used, except that the ORF50 deletion was introduced in the background of a kLANA-expressing MHV-68 mutant recently generated and kindly provided by Habison and colleagues [21]. Virus supernatants were produced similar to MHV-68Δ50. However, as we only achieved low titers with this construct we used higher volumes of the viral supernatants in our infections experiments (500 ul, diluted 1:2 in EBM-2 containing polybrene at a final concentration of 8 µg/ml polybrene, for infection of 5 x10^5^ MLE12 cells seeded into one well of a 6-well plate). The resulting cultures were sorted by FACS and either directly subjected to low cell ChIP, or were subcultured and sorted every 5 to 10 days by FACS to maintain a population of almost 100% infected / GFP-positive cells.

#### KSHV

Infectious wild-type KSHV supernatant was produced by inducing latently infected BCBL1 cells with TPA (Sigma-Aldrich; Cat#P8139-5MG) and sodium butyrate (Sigma-Aldrich; Cat#B5887-1G) followed by 100x concentration by centrifugation as described previously [4]. For infection, concentrated KSHV stock solutions were diluted in 600 µl EBM-2 medium without supplements (LONZA). Target cells were then incubated with virus for two hours followed by substitution with culture medium.

An ORF73 deletion mutant (KSHV-BAC16Δ73) of KSHV-BAC16 [46] was generated by en passant mutagenesis [47]. A PCR product consisting of ORF73 homology sites and a kanamycin resistance gene was generated to replace ORF73 in the KSHV-BAC16 by homologous recombination. In a second recombination step the kanamycin resistance gene was removed to obtain a deletion of nucleotides 140,572 to 143,943. The resulting KSHV-BAC16Δ73 was analyzed by restriction enzyme analysis with multiple restriction enzymes and high-throughput Illumina sequencing.

For the reconstitution of KSHV-BAC16Δ73 infectious viral particles, KSHV-BAC16Δ73 was transfected into iSLK cells [48] stably expressing kLANA with lipofectamine 2000 (ThermoFisher, Cat# 11668019). Two days post transfection, hygromycin selection (200 µg/ml) was initiated to select for bacmid containing cells (200 µg/ml). Virus production was performed as previously described [46]. For infection experiments employing KSHV-BAC16Δ73, 5 x 10^5^ MLE12-kLANA or MLE12-mLANA cells were seeded into one well of a 6-well plate, respectively. Cells were infected with 1 ml of virus supernatant of KSHV-BAC16Δ73 supplemented with 8 µg/ml polybrene. 2 hours post infection virus supernatant was removed and replaced by complete DMEM. Cells were subcultured in the presence of hygromycin (200 µg/ml) to maintain a homogenous population of 100% infected cells.

### Preparation of total RNA and protein extracts

Total RNA was extracted from MHV-68 infected cells using RNA-Bee (Tel-Test, Inc.; Cat#NC9850755) according to the manufacturer’s instructions. RIPA-extracts and Western Blot were performed using standard protocols.

### Antibodies

For ChIP the following antibodies were used in this study: Rabbit polyclonal anti H3K9/K14-ac (Merck Millipore; Cat#06-599); Rabbit monoclonal anti H3K4-me3 (clone MC315) (Merck Millipore; Cat#04-745); Rabbit polyclonal anti H3K9-me3 (Merck Millipore;Cat#17-625); Rabbit polyclonal H3K36me2 (MerckMillipore; Cat#07-274); Rabbit monoclonal H3K36me3 (Cell Signaling; Cat#4909S); Rabbit polyclonal JHDM1B/KDM2B (Merck Millipore; Cat#17-102-64, successfully tested in ChIP-seq: LOT 2135462, tested but non-functional in ChIP-seq: LOT 2756112 and LOT 3096002); Normal rabbit IgG (Merck Millipore;Cat#12-370). For detection of LANA expression the following antibodies were used: anti-kLANA rabbit polyclonal Ab, rabbit mLANA antiserum [49], and anti-ß-Actin mouse monoclonal Ab (Santa Cruz Biotechnology, Cat# sc-47778). The following secondary antibodies conjugated to horseradish peroxidase were used for western blot detection: Anti-rabbit IgG (Santa Cruz Biotechnology, Cat# sc-2004) and anti-mouse IgG (Invitrogen, Cat# 31431).

### Chromatin-immunoprecipitation (ChIP)

ChIP was performed essentially as described in detail previously [6]. Briefly, cells were treated with 1% formaldehyde in PBS for 10 min to crosslink proteins and DNA. Reactions were quenched by addition of glycine to a final concentration of 0.125 M. All buffers contained 1x protease inhibitor cocktail (Roche) and 1 mM PMFS. Cells were incubated in 1 ml buffer 1 (50 mM Hepes-KOH, 140 mM NaCl, 1 mM EDTA, 10% glycerol, 0.5% NP-40, 0.25% Triton X-100) for 10 min on ice to isolate nuclei. After centrifugation (1,350 x g 5 min), nuclei were washed with 1 ml buffer 2 (10 mM Tris-HCl, 200 mM NaCl, 1 mM EDTA, 0.5 mM EGTA). Pelleted nuclei were lysed in 1 ml buffer 3 (1% SDS, 10 mM EDTA, 50 mM Tris-HCl) by multiple pipetting and chromatin was fragmented to approximately nucleosome size using a BioruptorTM (Diagenode). After addition of 100 µl 10% Triton X-100 Cell debris was pelleted (20,000 x g, 4°C) and chromatin containing supernatant was collected. Chromatin of 1×10^6^ cells was diluted 1:10 in dilution buffer (0.01 % SDS, 1.1 % Triton X-100, 1.2 mM EDTA, 16.7 mM Tris-HCl, 167 mM NaCl). 10µg of the respective antibody was added to the sample and incubated for 16 hrs at 4°C rotating. 50 µl BSA-blocked ProteinG sepharose beads (GE Healthcare) were added to precipitate the chromatin-immunocomplexes and incubated for 1 hr at 4°C. Beads were washed once with 1ml of the following buffers: low-salt buffer (0.1 % SDS, 1 % Triton X-100, 2 mM EDTA, 20 mM Tris-HCl, 150 mM NaCl); high-salt buffer (0.1 % SDS, 1 % Triton X-100, 2 mM EDTA, 20 mM Tris-HCl, 500 mM NaCl); LiCl-wash buffer (0.25 M LiCl, 1 % Nonidet P-40, 1 % Na-deoxycholate, 1 mM EDTA, 10 mM Tris-HCl). Subsequently, beads were washed two times with TE buffer without protease inhibitors. Chromatin was eluted from the beads by incubation in 210 µl SDS containing elution-buffer (50 mM Tris-HCl pH 8.0, 10 mM EDTA, 1 % SDS) for 30 min at 65°C. Chromatin containing supernatant was separated from the beads by centrifugation. After addition of 8 µl of a 5 M NaCl stock solution chromatin was de-crosslinked at 65°C overnight. Contaminating RNA was degraded by addition of 200µl TE buffer containing 8 µl RNAseA (10 mg/ml) for 2 hrs at 37°C. Subsequently, 7 µl of CaCl_2_ solution (300 mM CaCl_2_ in 10 mM Tris-HCl) and 4 µl ProteinaseK (40 mg/ml) were added and incubated for 1h at 55°C to degrade proteins. DNA was purified by phenol-chloroform extraction and ethanol precipitation. Input DNA samples were diluted in 200µl elution buffer and treated identically to the ChIP samples beginning with overnight de-crosslinking. Finally, DNA was recoverd in 10 mM Tris-HCl and was either subjected to ChIP-seq library preparation or was analysed directly by ChIP-qPCR.

For Low cell ChIP we combined different protocols to facilitate efficient fragmentation of low numbers of cross-linked cells and library preparation from spurious amounts of DNA. Cell fixation was performed using the fixation reagents and steps from the Low Cell Chip Kit (Active Motif; Cat#53084) as described in the user manual. Samples were then treated with MNase according to the ‘MNase followed by sonication’ protocol described earlier [50]. Briefly, 200µl fixated samples containing 2000 to 9000 cells (depending on the sample) were diluted to a total volume of 200µl in dilution buffer supplemented with 3mM CaCl_2_. After incubation at 37°C for 15 minutes, MNase reaction was stopped by adding EDTA (10 mM final concentration) and EGTA (20 mM final concentration). Samples were sonicated using a Bioruptor similar to the ChIP protocol above for 10 cycles. After addition of Triton X-100 (0.1% final concentration) cell debris was removed via centrifugation. Total supernatant was collected and 400 µl were separated to serve as an input control. Chromatin precipitation was performed essentially as described in the ChIP section above only omitting the addition of CaCl_2_ during Proteinase K digestion of the input sample to prevent residual MNase activity therein. Sequencing libraries of low chip samples and input was performed using the library preparation reagents of the Low Cell Chip Kit.

### ChIP-qPCR

Relative enrichment of ChIP DNA was determined by qPCR and is presented as % of input. All primer used in this study are given in Table 1. Standard curves were generated for all primer sets to ensure efficient amplification. Specificity to the respective target site was assessed by melt curve analysis. All samples were analyzed on a Rotorgene 6000 qPCR machine (QIAgen) using the SensiMix™ SYBR® Hi-ROX Kit (Bioline, Cat#QT605-05).

**Table 1.**
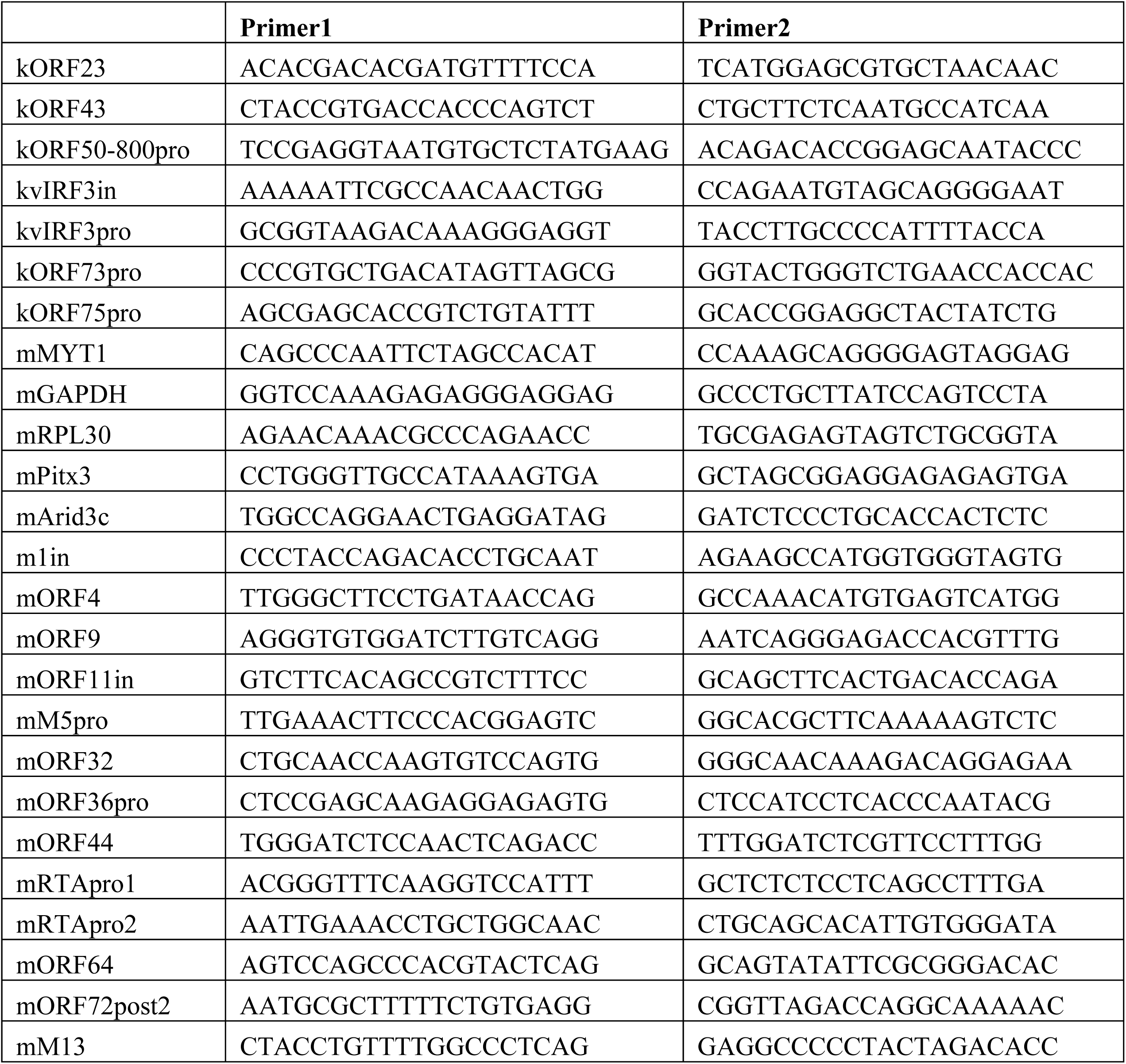
Primers used in this study.

### Library preparation and sequencing

ChIP and respective input libraries were prepared from 2-10 ng DNA using the NEXTflex Illumina ChIP-Seq Library Prep Kit (Bioo Scientific; Cat#5143-02) according to the manufactureŕs instructions. For RNA-seq, quality of extracted total RNA was verified prior to library preparation on a 2100 Bioanalyzer (Agilent). Strand specific Illumina compatible paired end ssRNA-seq libraries were generated using the NextFlex Directional RNA-Seq Kit (Bioo Scientific; Cat#5129-08) as per the manufacturer’s recommendations. Low cell RNA-seq libraries (unstranded) wer generated withSMART-Seq v4 Ultra Low Input RNA Kit for Sequencing (Takara/ Clonetech; Cat#634890). All sequencing libraries were sequenced on a HiSeq 2500 system (Illumina) using paired end (2×100bp) or single read (1×50) flow cells for RNA-seq and ChIP, respectively, or on a NextSeq 500 (Illumina) using single read (1×75) flow cells for ChIP and low cell RNA-seq.

### Sequencing data analysis

#### ChIP-seq

Quality filtered single end reads were aligned to the viral reference genomes of MHV-68 (NC_001826) and KSHV (HQ404500) as well as to the mouse (mm10) and human genomes (hg19) using Bowtie [51] with standard settings and the –m 1 option set to exclude multi mapping reads. Coverage calculation for visualization purpose was performed with IGV-Tools [52]. All coverage datasets used to generate graphs shown in Figs 1, 5, 6, 7, 8, 9, S2 Fig and S6 Fig are provided in S1 Dataset. For visualization purposes, ChIP-seq graphs were adjusted by subtracting the lower 5^th^ percentile to remove background signals [6]. Analysis and heatmap visualization of KDM2B ChIP-seq data as well as global H3K27-me3 and H3K36-me2 anti-correlation analysis was performed with EaSeq (http://easeq.net) [53].

#### Statistical analysis of histone modification and KDM2B enrichment on viral genomes

To calculate the relative enrichment of broad histone modifications H3K27-me3, H3K36-me2 and H2AK119-ubon viral genomes in comparison to the respective host genomes, we first detected all regions significantly enriched for the respective histone mark on the host genome against their input samples as controls using SICER/EPIC2 [54]. The 200 most significant (based on p-value) broadly enriched regions median sizes (15 to 40 kb for individual experiments) were selected as host positive regions and are named host pos in Figs 4, 5, 6, 7, 8 and in S7 Fig. The negative regions, which represent the general ChIP background of each individual analysis (host neg in the respective figures), were generated by random selection of regions with the same size distribution as the positive controls. We excluded SICER/EPIC2-positive regions as well as all regions with less than 10 reads per region in the input samples (the latter prevents selection of unmappable regions, e.g. in repetitive elements, in the negative controls) as well as all regions with known mappability bias (blacklists downloaded from ENCODE). Viral sequences were split into segments of 10kb (shifted by 5kb) to reflect broad regions. We counted the reads within each pos, neg and viral region using FeatureCounts. For each individual region we then calculated the ChIP to input read count ratio and normalized all groups to the median of the respective negative control. The resulting values represent relative enrichment of the respective broad histone modification signals over the general ChIP background present in each individual experiment. For quantification of relative KDM2B enrichment on the viral genome in comparison to host target loci (Fig 9), we detected KDM2B-bound sites using MACS peak calling similar to H3K4-me3 peak calling, selected control regions by FDR and performed the same analysis described above using 2kb windows (shifted by 1kb) for the KSHV genome.

#### Correlation coefficients

Correlation coefficients of ChIP tracks were generated using GraphPad Prism and are presented in S2 Dataset.

#### RNA-seq analysis

Quality filtered paired end reads were aligned to the MHV-68 sequence (NC_001826), to KSHV (HQ404500) as well as to the mouse genome (mm9) using the spliced read mapper STAR allowing for the detection of novel splice events in combination with FeatureCounts [55, 56]. For visualization purposes, coverage of both strands was calculated individually.

For ***expression analysis*** of MHV-68 encoded ORFs (Fig 2B and S2 Fig), all ORFs annotated in NC_001826 were assigned as single exons and genes in GFF3 format. Since the precise structure of mRNAs is not available for most ORFs, we evaluated coverage of coding regions to estimate viral transcription of individual gene products. Therefore, we restricted the analysis of viral gene expression by reads matching unique ORF regions only and ignored reads that could not be assigned to specific ORFs. Aligned reads were counted using FeatureCounts. Resulting feature counts of all viral ORFs in individual samples were normalized by the total number of reads that were aligned to the murine genome (mm9) by STAR. At later stages of lytic transcription, this normalization method may lead to an overestimation of general viral transcript levels but not to differences of the expression profile within each individual sample. Resulting data are given in S3 Dataset in raw and normalized format. Fig 2C depicts relative normalized expression data with the detection threshold set to a minimum of 10 reads per ORF.

### Statistical Information

For all figures containing statistical testing, we ensured that use of the respective test was justified as being appropriate and meeting the respective assumptions. For Figs 4, 5B, 6B, 7C, 8C, 9B and S7 Fig, data is presented as box-whisker-plot with 5th-95th percentile and mean (+). Significance was calculated by 1way ANOVA testing using GraphPad Prism. For Fig 5C, ChIP-qPCR data is presented as mean with SEM of three technical replicate ChIP-qPCR datasets normalized to the respective endogenous controls for H3K4-me3 and H3K27-me3 mGAPDH and mMYT1 using the primers given in Table 1. This ChIP-qPCR dataset originates from a third independent superinfection experiment that was performed to validate the two biological replicate ChIP-seq datasets of Fig 5B. For Fig 6C and 7A presented error bars represent SEM of at least two replicate experiments (for n see individual figure legends). For Fig 8A, ChIP-qPCR data is presented as mean with SEM of three technical replicate ChIP-qPCR experiments. The two independent datasets were generated from two mice that were de novo infected with MHV-68.

### Data Availability

All raw sequencing datasets used in this study will be made available via the European Nucleotide after peer review and formal acceptance of this article.

## Acknowledgements

We thank Kenneth M. Kaye and J. Pedro Simas for kindly providing the chimeric MHV68-kLANA bacmid, and Samuel H. Speck for kindly providing MHV-68-H2BYFP. We thank Daniela Indenbirken, Henry Scheibel, Kerstin Reumann and and Beatrix Steer for technical assistance, and Simon Weissmann and Nicole Fischer for helpful discussions and critical reading of the manuscript.

## Supporting Information

**S1 Fig.**
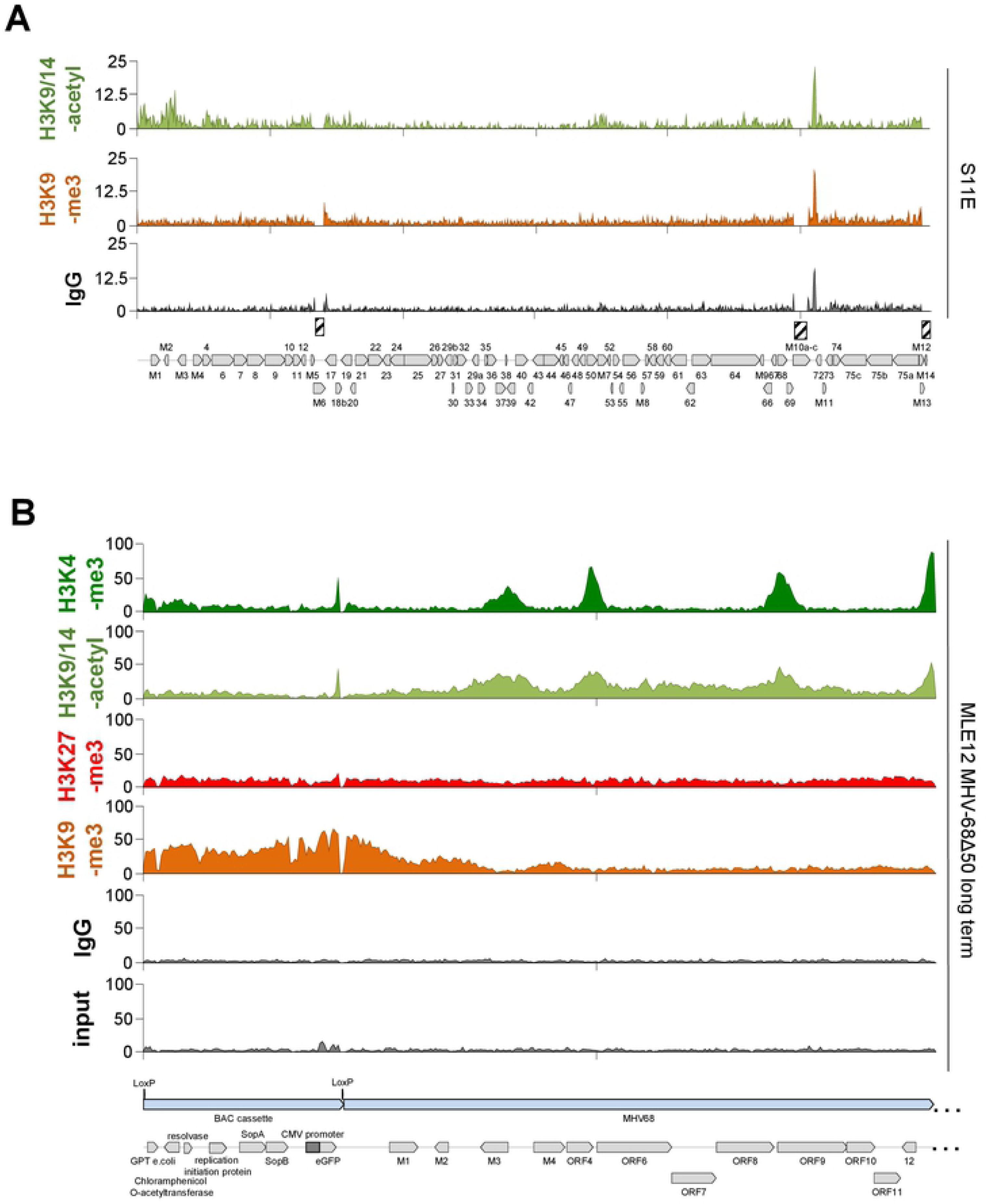
ChIP-seq coverage across the MHV-68 genome in S11E cells or the MHV-68Δ50 BAC cassette in MLE12 cells. Shown are coverage data from ChIP-seq experiments performed with the the indicated antibodies for (A) the MHV-68 genome in S11E cells or (B) the bacmid backbone at the leftmost end of the genome in MHV-68Δ50 BAC-infected MLE12 cells. For the latter, ChIP-seq data from the same samples shown in Figure 3B were mapped to the MHV-68 reference genome including the sequence of the BAC-cassette. For further details please refer to the legends of Figs 1 and 3.

**S2 Fig.**
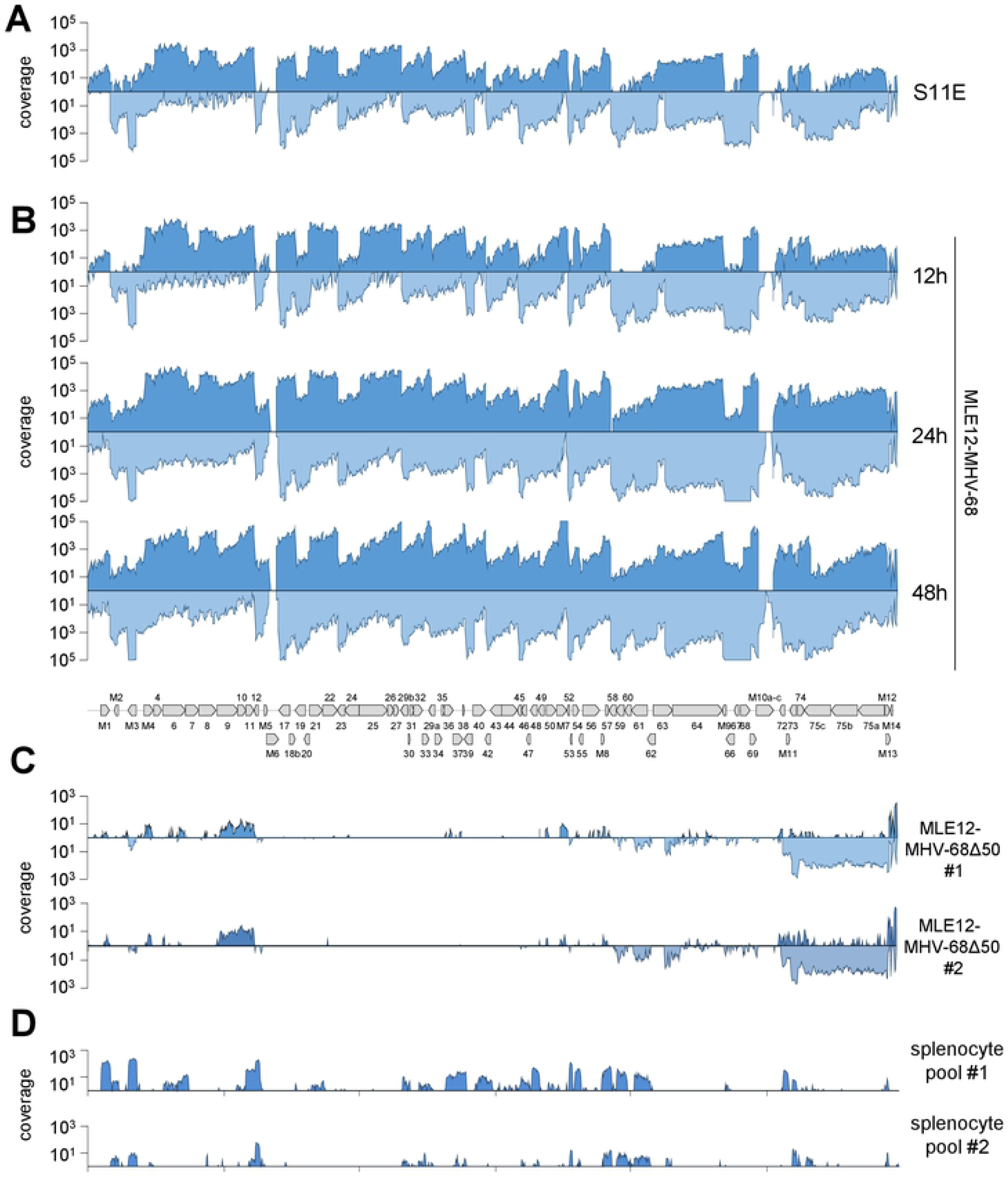
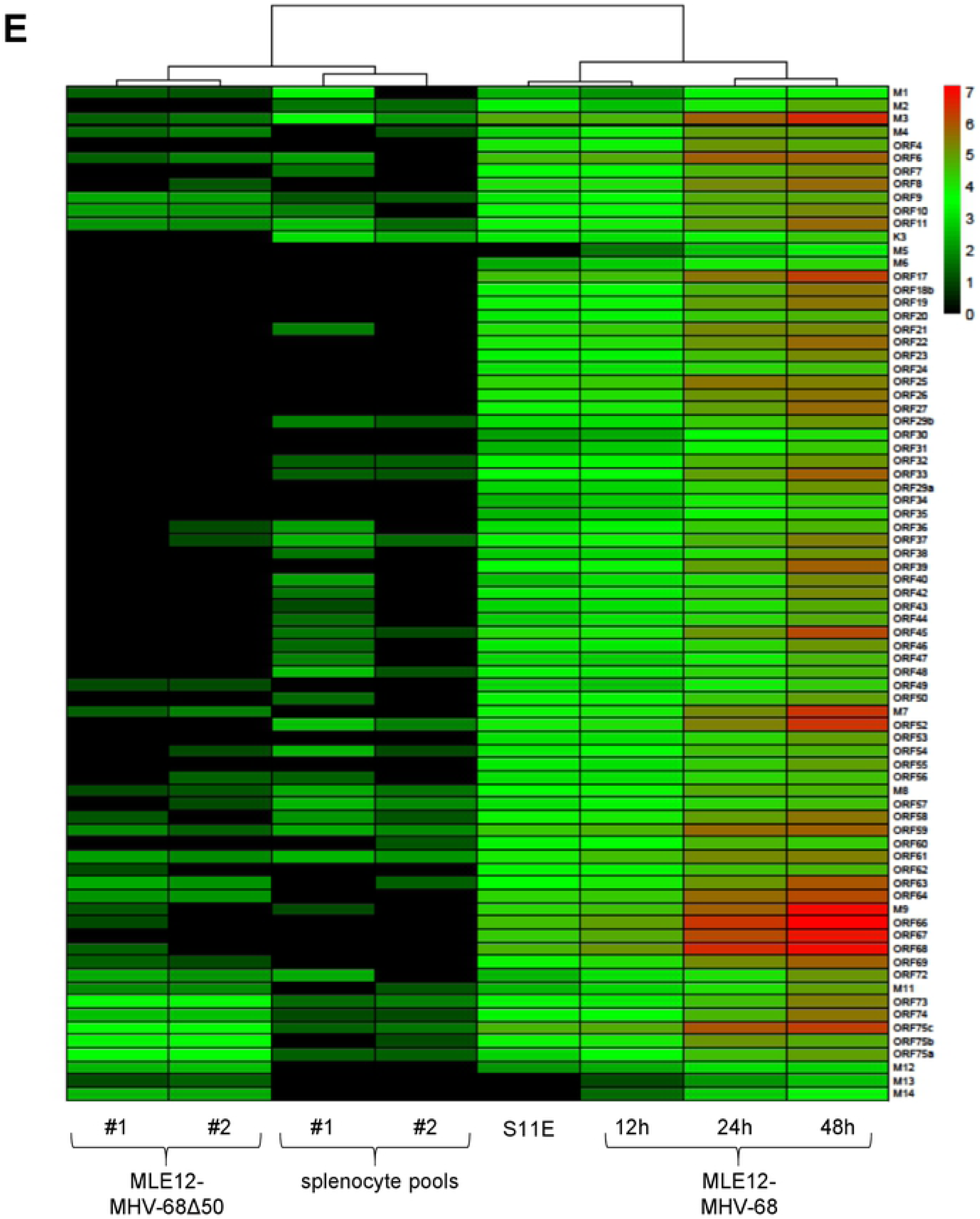
RNA-seq analysis of MHV-68 infected S11E and MLE12 cells. (A) RNA-seq analysis of persistently infected S11E cells (upper panel) or *de novo* MHV-68 infected MLE12 cells at 12, 24 and 48 hours post infection (lower panels). (B) RNA-seq analysis of two independent GFP-sorted MLE12 cell cultures which had been infected with MHV-68Δ50 for more than 3 weeks. (C) RNA-seq analysis of two splenocyte pools isolated from MHV-68-H2BYFP infected mice (3 mice per pool) at 17 days post infection. RNA sequencing for A and B was performed using a strand-specific sequencing protocol, for C a non-strand-specific, ultra-low input kit was used. Paired-end RNA-seq reads and single reads (for the low cell RNA-seq) were mapped to the MHV-68 reference sequence (NC_001826) using the splice-sensitive STAR pipeline (see Material and Methods for details). Coverage tracks depict mean coverage across 100 bp binning windows. For strand-specific data in A and B, forward and reverse strand coverage is shown in the upper and lower plots of each panel. Plots in C show coverage across both strands. (D) Heatmaps and hierarchal clustering (see tree at top) of normalized feature counts across individual MHV-68 ORFs annotated in the NC_001826 GenBank entry for the experiments shown in A, B and C.

**S3 Fig.**
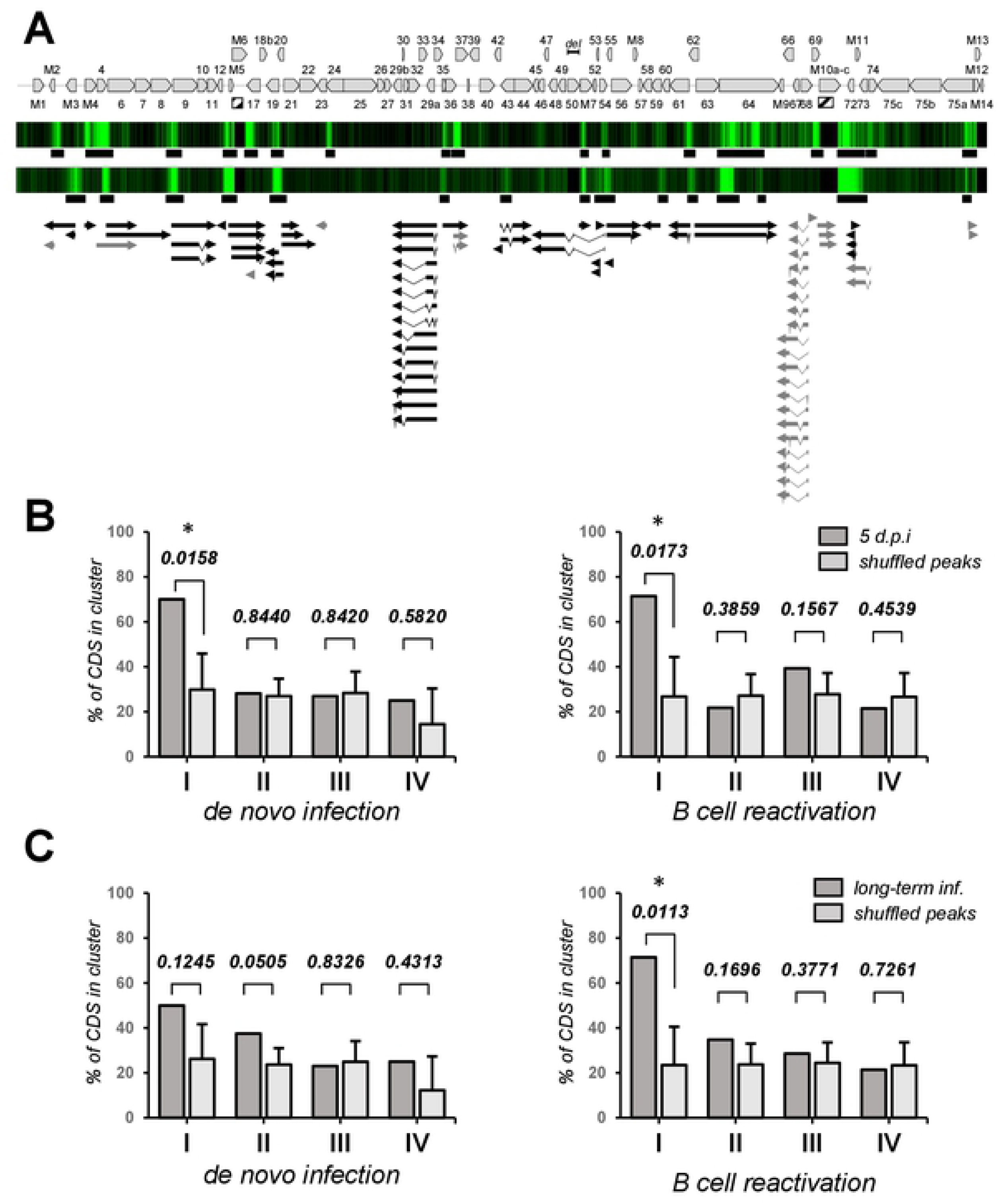
H3K4-me3 is enriched at putative mRNA start sites of immediate-early genes. (A) Black and dark grey arrows depict the predicted coding transcripts located downstream of an H3K4-me3 peak (as observed in MHV-68Δ50 infected MLE-12 cells) within a maximum distance of 250bp of their TSS. Transcripts downstream of peaks that are detected at 5 dpi but not in long-term infection are shown in gray. Tracks above transcripts reproduce the H3K4-me3 coverage from Fig 3 (top and bottom track correspond to data from 5 days p.i. or long term infected cultures, respectively) as a heat map, including the location of peaks detected by MACS14 (indicated by black bars underneath the tracks). (B, C) For each of the 4 expression kinetics clusters (I-IV) defined by Cheng and colleagues [57] for de novo infected fibroblasts (left graphs in each panel) or reactivated B-cells (right graphs) we calculated the percentage of ORFs encoded by transcripts located downstream of H3K4-me3 peaks observed after (B) 5 days of infection or in (C) long-term infected MLE-12 cells (dark grey columns in each graph). Light grey columns and associated error bars represent mean values and standard deviations of analyses repeated 100,000 times with randomly shuffled peaks. Cases with significant (<=0.05) p-values for the hypothesis that the number of ORFs observed with authentic peaks was significantly above that expected by chance (see S1 Protocol for further details) are indicated.

**S4 Fig.**
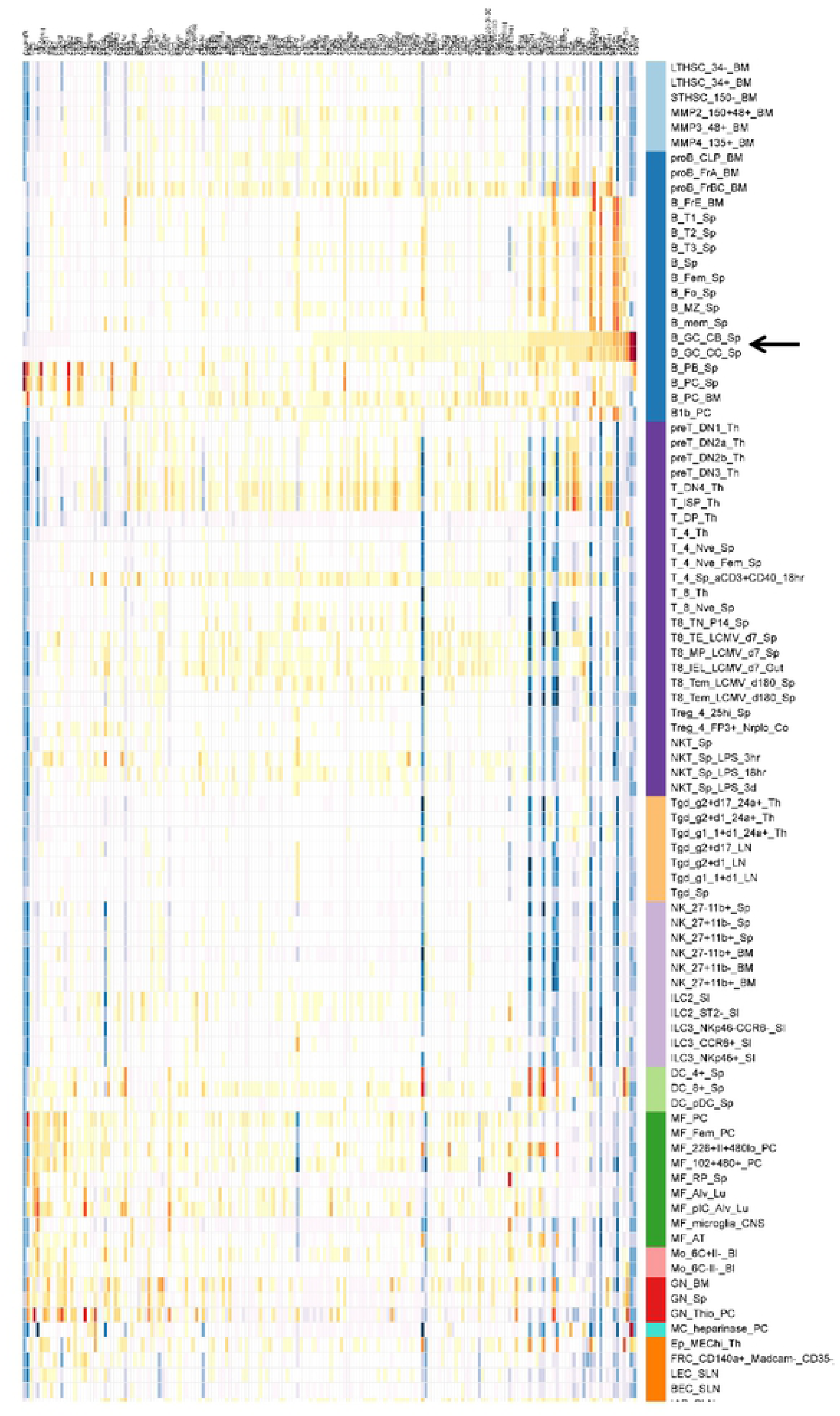
Immgen GeneSet analysis of 200 highly expressed genes in sorted infected B-cells. Immgen GeneSet analysis (http://www.immgen.org) of the top 200 expressed genes (as judged by STAR transcriptome analysis) from ultra-low input RNA-seq data of 1000 pooled splenocytes isolated from mice infected with MHV-68-H2BYFP 17 days post infection (see results and material & methods sections for details). The heatmap indicates the RNA-seq based row mean normalized expression values of the respective gene ID list for all immune cells within the Immgen database. Germinal center B-cells are indicated with an arrow.

**S5 Fig.**
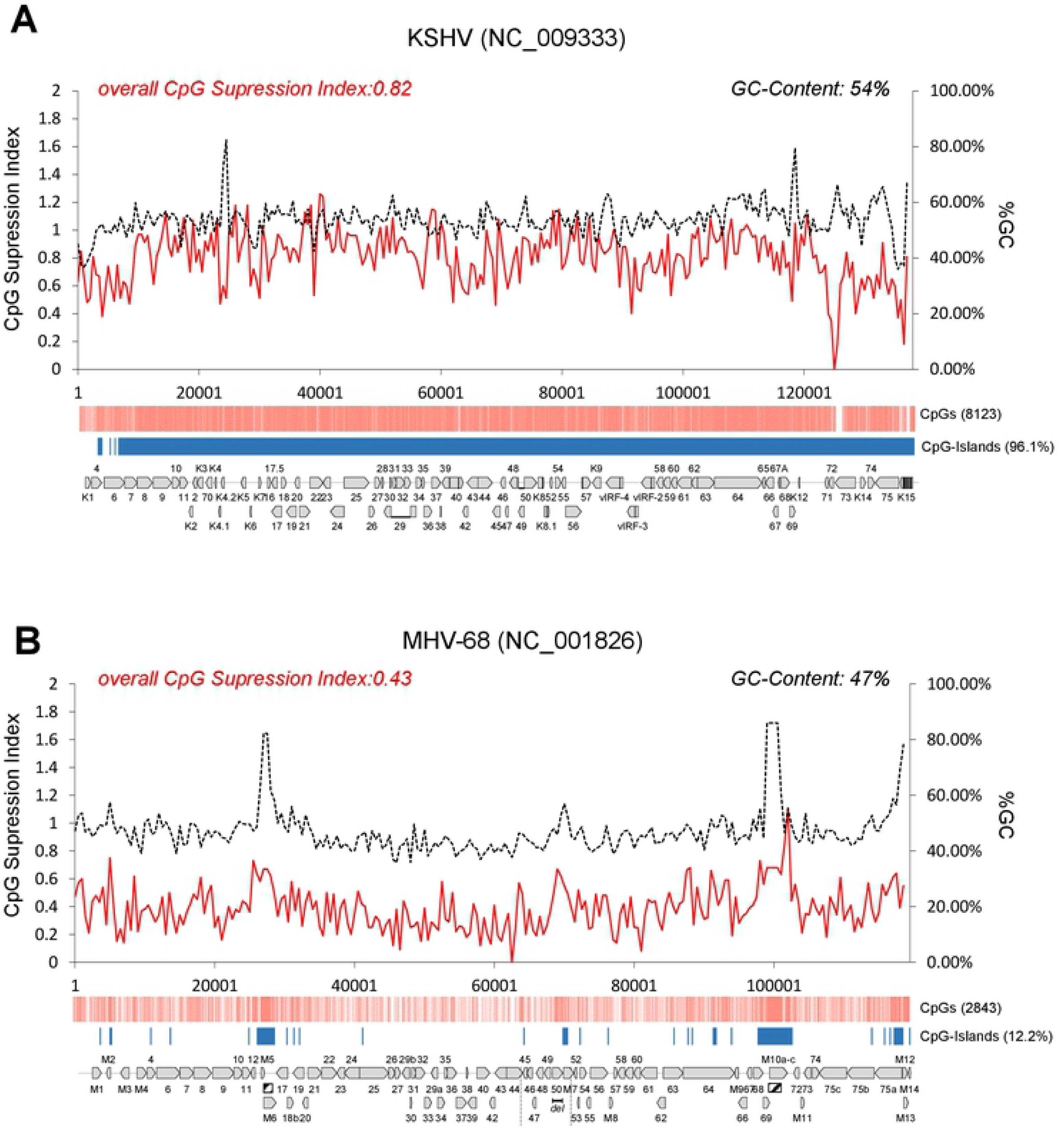
Analysis of CpG frequency/suppression and CpG island prediction in the genomes of KSHV and MHV-68. Graphs show GC content (black dashed line, right y-axis) and CpG supression index (red solid line, left y-axis) in a window of 500bp shifted in 250bp steps across the RefSeq genome sequences of (A) KSHV (GenBank accession NC_009333) or (B) MHV-68 (GenBank accession NC_001826). Overall CpG supression index and GC-content is indicated above the graph in each panel. The distribution of CpG motifs is shown in a map underneath the graphs, where the position of each individual motif is indicated by a vertical light-red line. The total number of CpG motifs is given to the right of the map. Blue bars below the CpG map indicate regions which register as CpG islands when employing the same criteria commonly used to designate host cell CpG islands (length >= 200bp, GC-content >= 50%, CpG suppression index >= 0.6). CpG islands were predicted by shifting a 200bp window in steps of 100bp across the viral genomes. Adjacent positive windows were then iteratively joined as long as the qualification criteria were satisfied by the merged region. The overall percentage of the viral genome that qualifies as a CpG Island is given to the right.

**S6 Fig.**
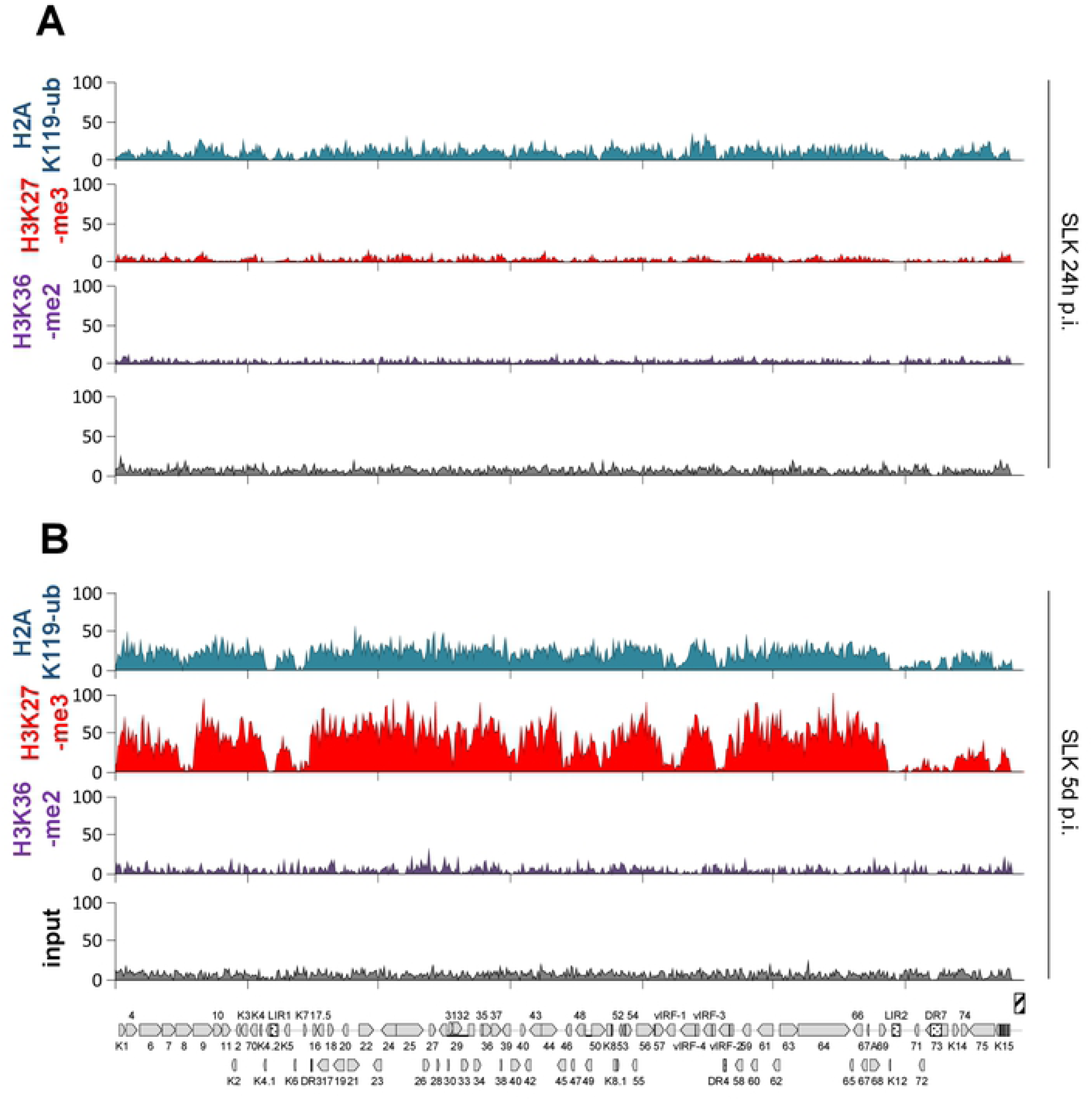
KSHV genomes acquire early H2AK199-ub marks upon infection of SLK cells. ChIP-seq coverage across the KSHV genome for (top) H2AK119-ub, (2nd from top) H3K27-m3, (2nd from bottom) or(bottom in each panel) input in SLK cells 24 hours (A) or 5 days (B) after infection with KSHV.

**S7 Fig.**
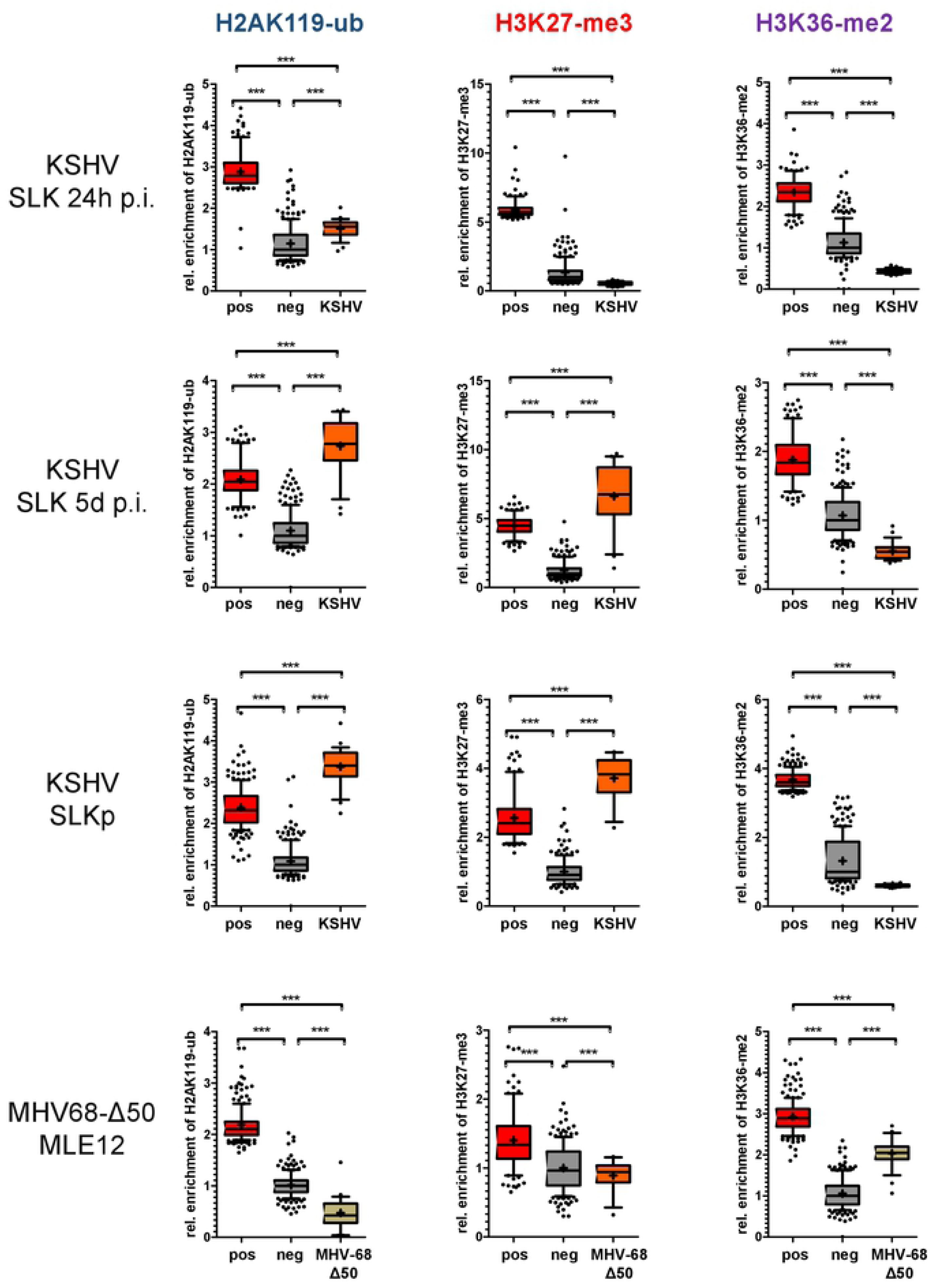
Statistical analysis of ChIP-seq data in de novo KSHV-infected SLK cells and MHV-68Δ50 infected MLE-12 cells. Enrichment of H2AK119-ub, H3K27-me3 and H3K36-me2 was analyzed using the statistical method described in the legend to Figure 4 and the materials and methods section. Data from MLE12 and SLKp cells correspond to those shown in Figs 9E and F for H2AK119-ub and H3K36-me2, or those in Fig 4 for H3K27-me3. Coverage tracks for SLK cells at 24 h.p.i and 5 d.p.i. are provided in S6 Fig.

**S8 Fig.**
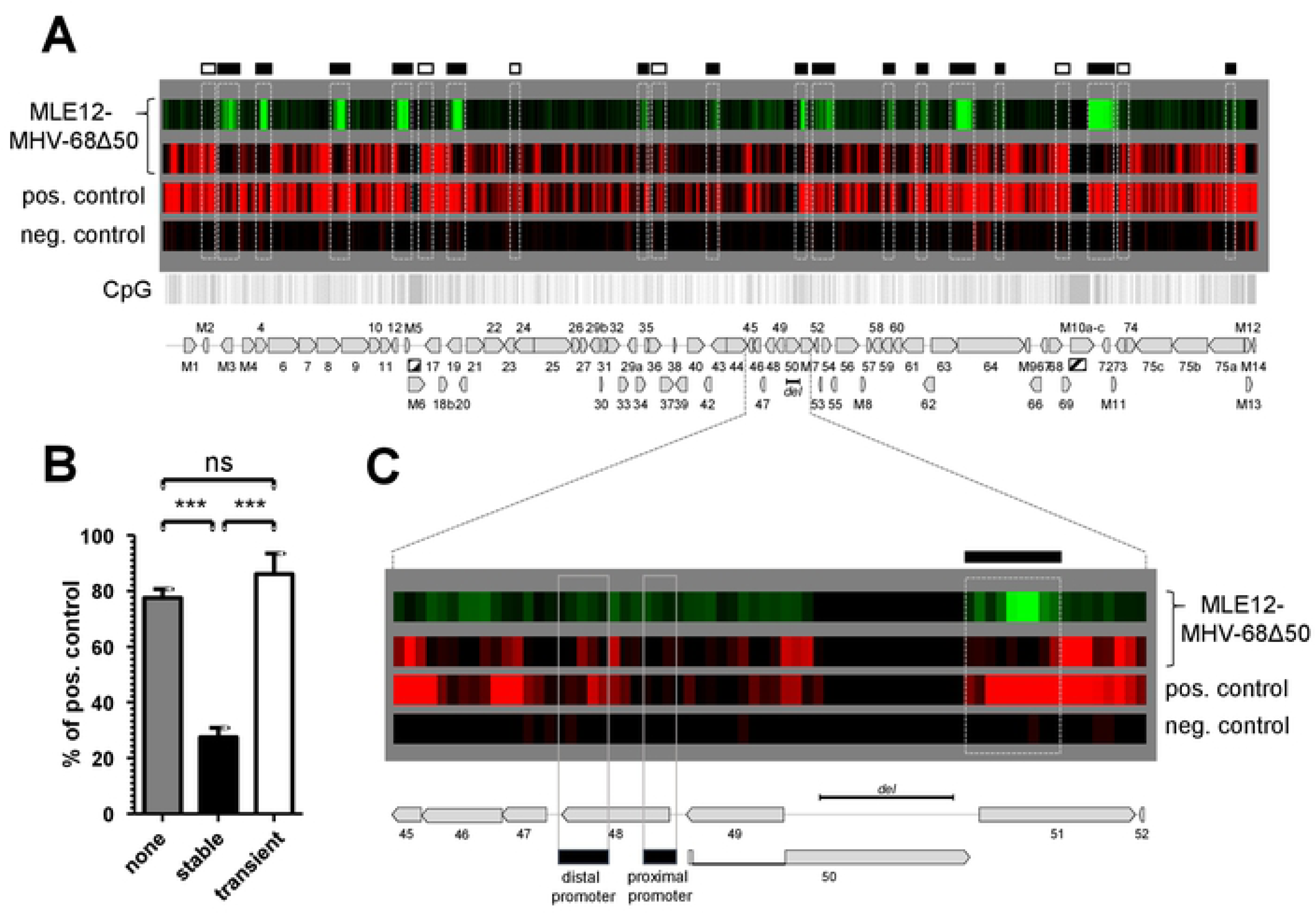
Analysis of CpG methylation by MeDIP-seq in long-term MHV-68Δ50-infected cells. DNA methylation levels were detected by MeDIP-seq using highly pure genomic DNA extracted from long-term MHV-68Δ50-infected MLE12 cells. We generated a positive control sample by in vitro methylation of MHV-68-BAC DNA that was spiked into genomic DNA from MLE12 cells to reflect authentic episome copy numbers in infected cells, as estimated by qPCR. Likewise, we used MLE12 DNA supplemented with unmethylated MHV-68-BAC DNA as negative control. Read coverage was normalized by total mapped read counts and viral input DNA to generate directly comparable tracks. (A) Episome-wide MeDIP-seq analysis. The three lower heat map tracks (in red) indicate relative MeDIP-seq coverage (normalized to the positive control) in the individual samples. The upper track (in green) reproduces the H3K4-me3 coverage data from Fig 3B. Boxes at the top of the panel indicate H3K4-me3 peaks detected by MACS in MHV-68Δ50-infected MLE12 cells. Filled boxes represent stable peaks which persist in long-term infected cells, whereas open boxes indicate transient peaks which are only observed at 5 days post infection. The peak positions are furthermore indicated by dashed frames overlaying the heat map panels. Hashed boxes in the genome map at the bottom indicate repetitive regions unsuitable for analysis and the dotted box marks the ORF50 deletion. (B) Quantification of relative DNA methylation levels (in percent of the positive control) from A in regions which do not acquire any H3K4-me3 peaks (none), or in stable or transient peaks (see legend to panel A for further information). Only persistent peaks protect viral DNA from methylation in long-term infected cells. Data are presented as mean ± SEM. (C) Detailed view of the ORF50 promoter region. The proximal and distal ORF50 promoters are marked by rectangles. The dotted box marks the deletion in the ORF50 coding region.

**S1 Dataset. ChIP-seq, RNA-seq and MeDIP-seq coverage tracks.**

This dataset contains all sequencing coverage tracks of MHV-68 and KSHV as presented in Figs 1, 2, 3, 5, 6, 7, 8, 9, S1, S2, and S6.

**S2 Dataset. Correlation coefficients.**

Correlation coefficients of all analyzed ChIP-seq data (S1 Dataset) were calculated using Graph Pad Prism.

**S3 Dataset. FeatureCounts data.**

This dataset contains raw and normalized results of STAR/FeatureCounts analysis of RNA-seq data. This data is presented in Fig 2C.

**S4 Dataset. H3K4-me3 peak data as detected by MACS14.**

**S5 Dataset. MHV-68 polyA-sites, splice-junctions, predicted primary transcripts / processed transcripts.**

This dataset provides all polyA-sites as well as splice junctions and frequencies observed in our RNA-seq analyses, along with a prediction of primary and processed transcripts. For the latter, the dataset further provides coding potential and proximity to H3K4-me3 peaks, as well as an estimate of relative abundance values in individual samples.

**S1 Protocol. Bioinformatic analysis of MHV-68 transcription units and correlation with activation-associated histone marks.**

**S2 Protocol. MeDIP-seq protocol.**

